# Strong postmating reproductive isolation in *Mimulus* section *Eunanus*

**DOI:** 10.1101/2022.12.21.521469

**Authors:** Matthew C. Farnitano, Andrea L. Sweigart

**Affiliations:** Department of Genetics, University of Georgia, Athens, GA, 30602, USA.

**Keywords:** postmating reproductive isolation, introgression, *Mimulus*, hybrid seed inviability, speciation

## Abstract

Postmating reproductive isolation can help maintain species boundaries when premating barriers to reproduction are incomplete. The strength and identity of postmating reproductive barriers are highly variable among diverging species, leading to questions about their genetic basis and evolutionary drivers. These questions have been tackled in model systems but are less often addressed with broader phylogenetic resolution. In this study we analyze patterns of genetic divergence alongside direct measures of postmating reproductive barriers in an overlooked group of sympatric species within the model monkeyflower genus, *Mimulus*. Within this *Mimulus brevipes* species group, we find substantial divergence among species, including a cryptic genetic lineage. However, rampant gene discordance and ancient signals of introgression suggest a complex history of divergence. In addition, we find multiple strong postmating barriers, including postmating prezygotic isolation, hybrid seed inviability, and hybrid male sterility, leading to complete or substantial postmating isolation in all species pairs. Hybrid seed inviability appears linked to differences in seed size, providing a window into possible developmental mechanisms underlying this reproductive barrier. While geographic proximity and incomplete mating isolation may have allowed gene flow within this group in the distant past, strong postmating reproductive barriers today are likely to prevent any ongoing hybridization. By producing foundational information about reproductive isolation and genomic divergence in this understudied group, we add new diversity and phylogenetic resolution to our understanding of the mechanisms of plant speciation.

## INTRODUCTION

Evolutionary biologists have long been interested in understanding what patterns and processes lead to reproductive isolation among species (Coyne & Orr, 1997; Darwin, 1859; V. Grant, 1981). Reproductive isolation is often broken down into components that act at sequential stages of the life cycle – premating barriers such as ecogeographic isolation and pollinator-mediated isolation that prevent mating, postmating prezygotic barriers that reduce fertilization success after mating has occurred, and postmating postzygotic barriers that reduce the fitness of hybrid offspring relative to pure species (Ramsey et al., 2003; Sobel & Chen, 2014). In plants, premating barriers are typically thought to evolve more quickly than postmating barriers and to have a greater contribution to overall isolation (Christie et al., 2022). However, this general pattern masks a great deal of heterogeneity across species pairs: many systems are completely isolated by premating barriers, while others rely exclusively on strong postmating isolation; many have a mix of both. Postmating barriers are often the result of intrinsic genetic incompatibilities, making them potentially more stable over the long term than ecologically mediated premating barriers (Coughlan & Matute, 2020). Postmating barriers may also be a source of cryptic variation in groups without obvious morphological or ecological differences.

To fully understand the role that postmating reproductive barriers play in speciation, they must be placed in a phylogenetic context. The degree of reproductive isolation between two taxa tends to increase with genetic distance (Christie & Strauss, 2018; Coyne & Orr, 1989; Malone & Fontenot, 2008; Scopece et al., 2007), but this relationship is not always consistent across taxa (Moyle et al., 2004) and is highly heterogeneous. In addition, individual reproductive barriers such as sterility and inviability may arise at different rates (Coyne & Orr, 1989; Le Gac et al., 2007; Malone & Fontenot, 2008; Presgraves, 2002). The reasons behind these differences are a subject of ongoing investigation, but may include differences in genetic architecture (Guerrero et al., 2017; Moyle & Payseur, 2009) or the forms of selection acting on each barrier (Baack et al., 2015). For example, conflicting selection pressures between maternal and paternal genomes in the mammalian placenta or the flowering plant endosperm are thought to give rise to reproductive barriers as these conflicts are resolved differently in independent lineages (Crespi & Nosil, 2013; Haig & Westoby, 1991; Lafon-Placette & Köhler, 2016); the relative strength of this parental conflict could influence the rate of barrier evolution (Coughlan et al., 2020; Raunsgard et al., 2018).

An excellent system for building a phylogenetically informed understanding of reproductive isolation is the western North American radiation of monkeyflowers in the genus *Mimulus.* Multiple species groups within *Mimulus* have already been studied extensively in a speciation context, including the *Mimulus guttatus* (e.g., Brandvain et al., 2014; Ferris et al., 2014; Lowry & Willis, 2010), *M. aurantiacus* (e.g., Stankowski et al., 2019; Streisfeld & Kohn, 2005), and *M. lewisii* (e.g., Nelson et al., 2021; Ramsey et al., 2003) species complexes. A number of pre- and postmating reproductive barriers have been mapped to single genes or QTL (e.g., Fishman et al., 2014; Fishman & Willis, 2006; Streisfeld & Rausher, 2009; Sweigart et al., 2006; Yuan et al., 2013; Zuellig & Sweigart, 2018). Recently, hybrid seed inviability in particular has been identified as a strong barrier in multiple independent species pairs of the *Mimulus guttatus* species complex, with parental conflict repeatedly implicated as a common mechanism (Coughlan et al., 2020; Oneal et al., 2016; Sandstedt et al., 2020; Sandstedt & Sweigart, 2022). An expanding trove of genetic and developmental resources across the genus allow for a holistic approach to understanding reproductive isolation from genic to phylogenetic scales (Yuan, 2019).

However, some *Mimulus* subgroups have received less attention, in part because they are less experimentally tractable or more difficult to access. Studies in these other groups would provide an important complement to existing knowledge, facilitating comparative analysis at the genus level while highlighting overlooked diversity in the genetic and ecological mechanisms of speciation. One such neglected group is *Mimulus* section *Eunanus* [synonym *Diplacus* section *Eunanus,* see (Barker et al., 2012; Lowry et al., 2019) for a discussion of nomenclatural issues]. Section *Eunanus* is a clade of ∼23 species (A. L. Grant, 1924; Nesom, 2013) which prior to this publication has had no genome-scale sequencing and few assessments of reproductive isolation. The group includes both widespread and narrowly endemic species occupying a range of elevations and habitats in western North America. Here, we focus primarily on three species from this group: *Mimulus brevipes* Benth.*, Mimulus fremontii* (Benth.) Gray*, and Mimulus johnstonii* Grant (Baldwin et al., 2012; A. L. Grant, 1924). These three morphologically defined species make up a well-supported clade according to a phylogenetic study of *Mimulus* based on three genetic markers (Beardsley et al., 2004). Their ranges are overlapping; *M. johnstonii* is restricted to higher elevations in the Transverse Ranges of southern California, while *M. fremontii* and *M. brevipes* are more widespread throughout the coastal ranges from Monterey into Baja California (Figure 1B).

**Figure 1.**
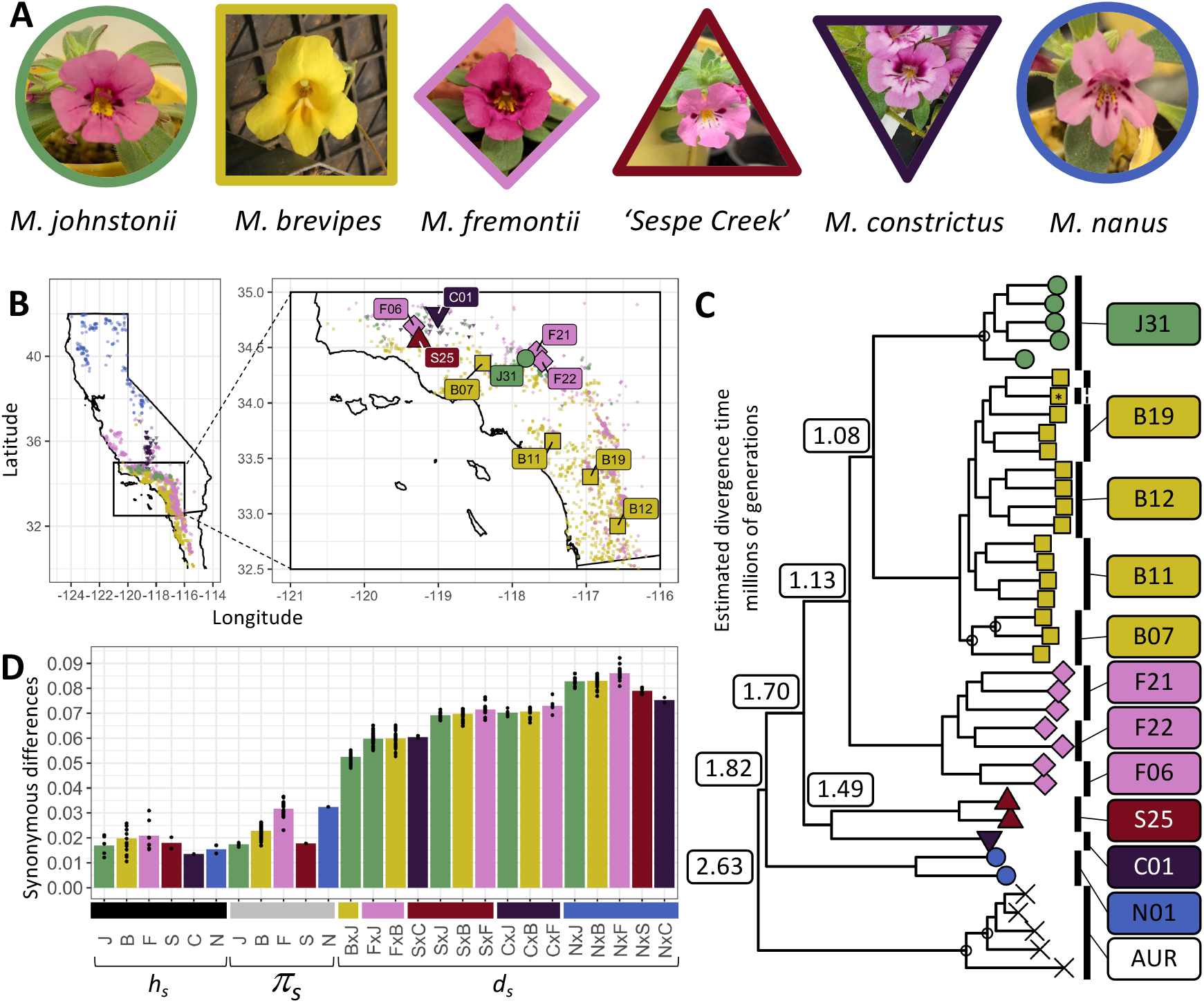
Phylogenetic relationships and divergence between sampled species in *Mimulus* section *Eunanus*. (A) Representative floral images for sampled species. (B) Map of occurrence records and sampled populations from the GBIF database for five species within California and Baja California. Inset highlights focal region for sampling, where species ranges overlap. Smaller icons indicate GBIF records (GBIF.org, 2022) for each species, which include iNaturalist research grade observations and herbarium collections. Larger named icons indicate sampled populations used in this study. The range of *M. nanus* extends into Nevada, Oregon, and Washington; population N01 (*M. nanus*) was collected from Washington state and is not shown. Population S25, referred to as Sespe Creek, is treated as a separate lineage. (C) Maximum likelihood phylogeny inferred using RAxML from the ‘complete’ dataset of genome- wide SNPs, rooted by the *M. aurantiacus* complex. All nodes except those marked with open circles have 100% bootstrap support. Species divergence times were estimated using synonymous divergence relative to synonymous diversity. All sampled populations are monophyletic with strong support, with the exception of one sample from B11 that clusters with B19 (marked with an asterisk). (D) Nucleotide heterozygosity (*h_s_*), pairwise diversity (*π_s_*), and divergence (*d_s_*) at genome-wide synonymous sites, indicating substantially higher between- species divergence than within-species diversity. Bars represent species or comparison means, while points represent individual pairwise sample comparisons. Diversity includes both within- and between-population comparison where possible. Sespe creek is substantially diverged from’

Studies of reproductive isolation in these three species are limited. Species distribution modeling between *Mimulus johnstonii* and *M. brevipes* identified strong but incomplete ecogeographical isolation (RI=0.65 and 0.82 reciprocally) (Sobel, 2014), and they can be found within hundreds of feet of each other (pers. obs.). No distribution models have been made with *M. fremontii*, but M*. brevipes* and *M. fremontii* are found at similar elevations, flower at similar times, and can be found growing within inches of each other (pers. obs.). Major differences in flower color, size, and shape between *M. brevipes* and its relatives might suggest some degree of pollinator-mediated isolation, though little is known about the identity of pollinators or levels of outcrossing; *M. fremontii* and *M. johnstonii* have similar floral characteristics and seem unlikely to have substantial pollinator isolation. Postmating barriers have not been studied, with the exception of one study showing low germination rates of F1 hybrid seeds between *M. brevipes* and *M. johnstonii,* potentially suggesting hybrid seed inviability (Sobel, 2010).

In this study, we present the first genome-scale analysis of divergence relationships in *Mimulus* section *Eunanus*, incorporating our three focal species as well as two more distant species, *M. constrictus* and *M. nanus*. In addition, we use controlled crosses to examine multiple postmating reproductive barriers between the focal species. We find clear genetic divergence and multiple strong postmating reproductive barriers between all tested species. Our results add to a growing body of evidence that intrinsic postmating barriers, especially hybrid seed inviability, are a key component in the maintenance of plant species.

## MATERIALS AND METHODS

### Sample collections and growth conditions

Fruits were collected from wild plants of three focal species (*M. brevipes, M. fremontii,* and *M. johnstonii*) at nine locations in southern California in summer 2019 (Figure 1B, Table S1). As additional, more phylogenetically distant comparisons, a population of *M. nanus* in Oregon and a population of *M. constrictus* from California were collected in 2021. To germinate, we treated seeds overnight with 1 mM giberrellic acid, then rinsed three times with distilled water and placed them individually in 2x2cm soil plugs. Soil was a 1:1:1 mix of sand, perlite, and peat with a thin layer of finely sifted soil mix on top. Germination trays were set in standing bottom water and kept moist with a squirt bottle as necessary. When germination rates were low, liquid smoke was added to standing water to simulate post-fire conditions and stimulate germination. After germinants developed one or two pairs of true leaves, we transplanted soil plugs into plastic conical pots filled with the same soil mix. The conical pots were maintained in standing bottom water in a growth chamber set to 23C days/18C nights, with low relative humidity and 16 hours of daylight. Germination and establishment was low overall and highly variable across species, populations, and trials, so germination and establishment rates were not used as a proxy for seed viability or hybrid fitness.

One population initially collected as *M. johnstonii,* S25 from Sespe Creek, showed morphological differences suggesting it may be a unique lineage, and is treated separately throughout this paper. Compared to *M. johnstonii*, Sespe Creek has inserted stigmas, a lighter purple corolla with a slight white wash at the mouth, and longer leaf trichomes.

### Genomic sequencing

We collected leaf tissue from 33 individuals for genomic sequencing. All sequenced individuals were generated from outbred field-collected seeds and germinated in growth chambers. We flash-froze leaf tissue in liquid nitrogen and used a modified CTAB-based protocol to extract genomic DNA (Fishman, 2020). Briefly, tissue was ground to a powder and incubated in CTAB buffer with ß-mercaptoethanol. Supernatant was extracted with phenol-chloroform-isoamyl alcohol, then again with chloroform-isoamyl alcohol, followed by precipitation of carbohydrates with sodium chloride and polyethylene glycol. DNA was then precipitated with isopropanol, washed with ethanol, and resuspended in distilled water.

We used the tagmentation-based Illumina Nextera XT DNA library prep kit (FC-131-1024) to prepare genomic DNA for sequencing. The 33 individually barcoded samples were then sequenced in four batches at the Duke University Sequencing and Genomic Technologies Shared Resource with either an Illumina NextSeq 500 or an Illumina NovaSeq 6000 sequencer, in combined lanes with other barcoded projects (Table S2).

### Sequence alignments and SNP calling

We combined our dataset of 33 samples with five previously published samples (Stankowski et al., 2019) from the *Mimulus aurantiacus* species complex as an outgroup for analysis. *M. aurantiacus* is outside of section *Eunanus* but is the closest group with previous genome-scale data, as well as the closest group for which a reference genome assembly is currently available. For all 38 samples, we trimmed raw sequencing reads using Trimmomatic v0.39 (Bolger et al., 2014) to remove adapters and low-quality ends. We mapped reads to the *Mimulus aurantiacus* reference genome (Stankowski et al., 2019) using bwa mem v0.7.17 (Li & Durbin, 2009), then sorted and indexed with samtools v1.10 (Danecek et al., 2021). We removed PCR duplicates using the MarkDuplicates tool in Picard v2.21.6 (Broad Institute, 2019), and removed reads with unmapped mate pairs or mapping quality <29 using samtools view (Danecek et al., 2021). Average read coverage per sample was 2.2-15.7 (mean 6.8) across 33 novel samples (details in Table S1).

We called SNPs in each sample using the GATK v4.1.6.0 haplotype caller, then used GATK GenotypeGVCFs to jointly call variant and invariant sites for the entire dataset in ‘all-sites’ mode (Van der Auwera & O’Connor, 2020). Only sites mapping to the ten major linkage groups of the reference genome, corresponding to ten nuclear chromosomes, were included. We split the resulting VCF into SNPs and invariant sites, and separately filtered each before recombining them into a joint dataset, following recommendations by Dmitri Kryvokhyza (Kryvokhyzha, 2022). We filtered both SNPs and invariant sites to remove sites with combined DP <83 (∼1/3 of mean combined depth) or >1376 (two standard deviations above mean combined depth), QD<2, SOR>3, or MQ<40. We additionally filtered SNPs to keep only biallelic sites across the entire dataset, and to remove sites with QUAL<40, FS>60, MQRankSum<-12.5, ReadPosRankSum>12.5, or ReadPosRankSum<-12.5.

Starting with all sites that pass the above filters (14,814,282 biallelic SNPs and 96,871,941 invariant sites), we generated four datasets for analysis. For genome-wide phylogenetic and introgression analyses, we used a ‘complete’ dataset, which retained only sites where fewer than 30 of 38 (approximately 80%) of samples had a called genotype, resulting in 7,728,322 biallelic SNPs and 32,658,171 invariant sites. When calculating genome- wide introgression metrics, we also attempted to reduce stemming from uneven sampling of species and uneven coverage across individuals, by creating a ‘downsampled’ dataset of equalized sampling and coverage. For this set, we chose the two most divergent individuals from each species (except for *M. constrictus* for which we only had one sample), then randomly sampled 12 million raw read pairs from each individual and used the same pipeline as above to generate SNPs and invariant sites. Again, we retained only sites where 10 of 13 samples (∼80%) had called genotypes, resulting in 3,778,609 SNPs and 26,939,196 invariant sites.

For the calculation of diversity and divergence metrics, we made a ‘synonymous sites’ dataset. Starting with the ‘complete’ dataset, we used the *M. aurantiacus* gene annotation to select only sites in the 3^rd^ codon position of coding sequences that were four-fold degenerate, i.e., any nucleotide change to that site in the reference background would not change the protein sequence, using a custom script by Tim Sackton (Sackton, 2014/2022). The ‘synonymous sites’ dataset included 967,470 SNPs and 1,654,798 invariant sites. We note that multiple mutation hits within a single codon are possible given the large proportion of variant sites across the dataset, which could change the synonymous nature of these sites, but we expect these events to be rare enough to not substantially affect our genome-wide results on average.

For gene-by-gene analyses, we used a ‘genic’ dataset that included all sites passing the initial filters that fell within each of the 22,421 genes in the *M. aurantiacus* genome annotation. *Phylogenetic tree building*

To determine phylogenetic relationships among our samples, we built a maximum likelihood phylogenetic tree using the GTR+Gamma model in RAxML v8.2.12 (Stamatakis, 2014) with SNPs from the ‘complete’ dataset. SNPs were extracted into a concatenated fasta alignment using the ‘consensus’ function from bcftools v1.13 (Danecek et al., 2021) and invariant sites were excluded. Heterozygous sites were initially coded as IUPAC ambiguity codes, then randomly converted to a single allele with the ‘randbase’ function in seqtk v1.3 (Li, 2018). We ran 10 separate iterations of RAxML with unique random seeds, then chose the iteration with the largest log-likelihood. We used the RAxML -b option to run 1000 rapid bootstrap trees. We also generated a neighbor-joining tree with 1000 bootstrap replicates from the same data, using the dist.ml, nj, and boostrap.phyDat functions in the R package phangorn (Schliep, 2011).

We generated gene trees to examine patterns of gene tree discordance on the phylogeny. We used SNPs from the ‘genic’ dataset in the *M. aurantiacus* genome annotation, following the same approach as above to generate a maximum-likelihood phylogenetic tree in RAxML for each gene. Genes with insufficient data to resolve a tree (any individual with no called sites, or any two individuals with identical sequence) were excluded, resulting in trees for 17,573 genes. We then input these trees into ASTRAL v5.6.1 (Rabiee et al., 2019) which uses a multispecies coalescent approach to calculate a consensus species tree and quartet support scores for each node.

### Quantifying homozygosity, diversity, and divergence

Pairwise sequence differences were calculated from the ‘synonymous sites’ dataset using the ‘dxy’ and ‘pi’ functions in the program pixy v1.2.3 (Korunes & Samuk, 2021). For each comparison, sequence differences were calculated in 1Mb windows across the genome, then added together and divided by the total number of informative sites. We calculated pairwise diversity (π_s_) for each pair of individuals within a species, pairwise divergence (*d_s_*) for each pair of individuals across species, and heterozygosity (*h_s_,* pairwise diversity within an individual) for each individual. Diversity, divergence, and heterozygosity values were averaged across individuals within each species, or across individual pairs within each species pair.

To estimate the divergence times between species, we used the following molecular clock formula: (divergence time in generations) = (synonymous divergence – synonymous ancestral diversity)/(2*mutation rate). We assumed ancestral diversity for any two species to be the average of within-species diversity for the two species, and took the mutation rate to be 1.5x10^-8 (Koch et al., 2000). These estimates are approximate, as they do not take into account the effect of changing effective population sizes, such as demographic bottlenecks, or the possibility that ancestral diversity was very different from current diversity.

### Quantifying introgression

To explore historical introgression between species, we used three complementary approaches: ABBA-BABA tests and the related *f-*statistics; gene tree discordance bias using TWISST; and the model-based TreeMix program.

We used Dsuite v0.4 (Malinsky et al., 2021) to calculate Patterson’s D, a measure of the relative frequency of ABBA and BABA sites in the genome, and the related *f4* statistic, which uses allele frequencies to estimate the proportion of the genome resulting from introgression. We calculated D and *f4* for all possible trios of species in our dataset, using both the ‘complete’ and ‘downsampled’ SNP datasets, with the *M. aurantiacus* complex as the outgroup. Z-scores and associated p-values were calculated for each trio using a block-jackknife approach with 100 blocks in Dsuite; to account for multiple tests, a Bonferroni correction was used to obtain a corrected p-value (p_corr_). Because there are multiple trios available to estimate introgression for the same pair of species, we summarized results across trios using the *F_branch_* statistic in Dsuite for each species pair (Malinsky *et al*. 2018).

We used TWISST (github version d56cefb) (Martin & Van Belleghem, 2017) to obtain the proportion of gene trees supporting particular tree topologies. For each trio of species, we compared the two possible alternate (different from the consensus specie tree) tree topologies using a binomial test with expected ratio 1:1 to look for an excess of one topology over another. The *M. aurantiacus* complex was always used as the outgroup. This test is analogous to the SNP-based ABBA-BABA test, but uses gene trees rather than SNPs.

We used TreeMix (Pickrell & Pritchard, 2012) to create phylogenetic network models with or without migration edges as a further test of introgression. We used the ‘synonymous sites’ dataset, including only variant SNPs with no missing data, and grouped individuals by species. The dataset was pruned to select SNPs not in linkage disequilibrium using a custom script by Joana Meier (https://github.com/joanam/scripts/raw/master/ldPruning.sh). We ran models with zero to four migration edges, with 10 replicate runs per model type. The program OptM (Fitak, 2021) was used to define the best fitting TreeMix migration model.

### Identifying candidate windows with historical introgression

To identify candidate genomic regions with signatures of historical introgression, we used Dsuite (Malinsky et al., 2021) to calculate the window-based *df* statistic (Pfeifer and Kapan, 2019) for windows of 100 informative SNPs, each overlapping the previous by 50 SNPs. We calculated *df* across the genome for three focal trios with strong support for introgression, using the *M. aurantiacus* complex as the outgroup. In each quartet, we selected the 100 windows (out of 9594 to 17530, depending on the quartet) with the most extreme (positive or negative) values as potential candidates of introgression, with positive values set to match the direction of the genome-wide *F_branch_* signal.

### Measuring reproductive barriers

We assessed three different barriers to reproduction: postmating prezygotic isolation, F1 hybrid seed inviability, and F1 hybrid sterility. We conducted hand pollinations of greenhouse-grown plants within and among species by first removing anthers from the maternal flower to prevent self-pollination, then using forceps to place pollen from the paternal flower on the maternal stigma surface. Stigma lobes in these species close quickly in response to touch, so hand-pollination is not always successful. In a few cases, supplemental pollen from the same paternal plant was added the following day once the stigma had re-opened. We collected mature, browned fruits, categorized according to cross type. Cross type throughout this paper refers to the combination of maternal species and paternal species, with maternal species always listed first; the 16 cross types are JxJ, JxB, BxJ, JxF, FxJ, JxS, SxJ, BxB, BxF, FxB, BxS, SxB, FxF, FxS, SxF, SxS where J=*M. johnstonii*, B=*M. brevipes,* F=*M. fremontii*, S=Sespe creek population. For the cross type BxF, we also germinated and grew F1 hybrids from four independent crosses in the same conditions as parental plants (Table S3). These F1 hybrids were crossed to themselves and reciprocally with *M. fremontii* and *M. brevipes* as above (cross types HxB, BxH, HxH, HxF, FxH, where H=F1 hybrid). F1 hybrids between other species pairs were not grown because of low crossing success or low seed viability.

To measure postmating prezygotic isolation, we quantified crossing success (probability that a pollination produces at least one seed) and seed production (number of seeds per fruit) after hand-pollination; both viable and inviable seeds were included but unfertilized ovules were not. To test for an association between cross type and crossing success, we ran a penalized Firth regression with a binomial family function using the R package ‘brglm’. A Firth regression was chosen to account for complete separation (all successes or all failures) of some groups. To test the dominance relationships of this barrier, we compared the crossing success of *M. brevipes* x *M. fremontii* F1 hybrids to their parental species. We ran a generalized linear mixed-effects model (GLMM) with a binomial family and logit link function using the R package ‘lme4’ on cross types BxB, BxH, BxF, HxB, HxH, HxF, FxB, FxH, and FxF, where H indicates F1 hybrid offspring from a BxF cross. For this model, maternal population was included as a random effect, treating each independent F1 cross as its own population.

To assess seed viability in each cross, we used a combination of visual inspection and chemical treatment. All seeds were inspected under a dissecting scope and scored as inviable or viable. Viable seeds were plump and generally ovate, although they were allowed to be slightly misshapen as long as they appeared to be full. Inviable seeds included any seed that was severely shriveled, concave, empty, much smaller than the typical within-species seed, or with a shape very different from the typical ovate shape. To confirm that seeds visually scored as inviable were in fact inviable, we treated a subset of seeds with tetrazolium chloride, which stains viable seeds dark red. For this treatment, seeds were scarified using 1:5 bleach:distilled water with 0.83 uL/mL Triton-X for 15 minutes, washed twice with distilled water, and placed in 1% (w/v) tetrazolium chloride, then incubated in the dark at room temperature for ∼48 hours.

To test for an association between cross type and the relative counts of viable vs. inviable seeds, we ran a penalized Firth regression using the R package ‘brglm’ with a binomial family and logit link function. A Firth regression was chosen to account for complete separation (all viable or all inviable) of some groups.

To investigate reciprocal differences in seed size, a predicted correlate of parental conflict-driven hybrid seed inviability (Haig & Westoby, 1991), we imaged a subset of seeds from 49 fruits under a dissecting scope and used imageJ to manually measure seed length and width. This included all available seeds for those cross types with few seeds, and a random selection of seeds from a random selection of fruits for other cross types. We ran a linear mixed-effects model using the R package ‘lme4’ to test for differences in seed length between the four intraspecific cross types (JxJ, BxB, FxF, and SxS), using fruit as a random effect. We also ran six independent LMMs, one for each pair of species, to directly compare intra- and interspecific seeds. For example, we tested the *M. johnstonii* vs. *M. brevipes* pair by comparing cross types JxJ, JxB, BxJ, BxB. For each species-pair, we ran one LMM to test for an association between cross type and seed length, and a second to test for an association between cross type and seed length/width ratio.

To assess hybrid male sterility of *M. brevipes-M. fremontii* F1 plants compared with their parental species, we scored pollen number and pollen viability using an aniline blue stain. For each individual, all anthers from a single flower were collected in 50 µL 0.25% aniline blue in lactophenol solution. Blue stained pollen grains were scored as viable, while unstained clear pollen grains were counted as inviable. Pollen was counted in full 1mm^2^ squares of a hemocytometer until 9 squares or at least 100 total pollen grains were counted. For each individual, we estimated pollen viability by determining the proportion of viable pollen grains out of the total counted (for some individuals, pollen viability was measured from multiple flowers and the average was used). We used an LMM to test for an association between species ID and the total count of pollen grains scored per hemocytometer square, with population as a random effect. Similarly, we used a GLM with a binomial family and logit link function to test for an association between species ID (*M. brevipes, M. fremontii,* or F1 hybrid) and the relative counts of viable vs. inviable pollen grains, adding population as a random effect (each hybrid family was treated as its own population).

To assess *M. brevipes-M. fremontii* F1 female fertility, we hand-pollinated F1 hybrids with pollen from either parent and measured seed production. To determine whether F1 hybrids produce fewer seeds than within-species crosses, we used a linear model to test for an association between cross type and the number of seeds produced per fruit, comparing cross types BxB, BxH, HxF, and FxF and only counting fruits that produced at least one seed.

For every statistical model above, we ran Tukey post-hoc tests implemented in the package ‘multcomp’ to test for pairwise differences between each cross type or group.

We followed the methods of Sobel and Chen (2014) to calculate standardized measures of reproductive isolation for each of three reproductive barriers, as well as total postmating reproductive isolation, between each species. All measures used the equation

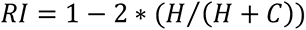

where H=heterospecific fitness and C=conspecific fitness. In this framework, RI ranges from -1 to 1 where -1 is complete heterosis, 0 is random assortment, and 1 is complete isolation. We calculated total measured reproductive isolation sequentially with the equation

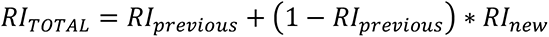

where RIprevious is the total RI from all previous barriers and RInew is the next barrier. Barriers that could not be measured due to complete or near-complete previous RI were left blank.

## RESULTS

### Genome-wide variation in the *M. brevipes* group defines clear species but also reveals a complex history of introgression.

A maximum-likelihood phylogeny generated from whole-genome SNP data resolves each of the three focal species (*M. brevipes*, *M. johnstonii*, and *M. fremontii*) as monophyletic with 100% bootstrap support (Figure 1C). Within these species, each population is also recovered as monophyletic, with the exception of one sample from a *M. brevipes* population (Figure 1C: one individual from population B11 clusters with B19). *M. johnstonii* is supported as sister to *M. brevipes,* in contrast to (Beardsley et al., 2004) which found *M. brevipes* and *M. fremontii* as sister based on just three loci. Genome-wide estimates of pairwise nucleotide divergence at synonymous sites also support strong divergence among the three focal species, with pairwise divergence values (*d_s_*, 4.8-6.0%) consistently well outside the range of nucleotide diversity within species (*π_s_*, range 1.6-3.7%) (Figure 1D, Table S4). Heterozygosity is similar to pairwise diversity (*h_s_*, range 1.1-3.1%), supporting a primarily outcrossing strategy in these species. Speciation times were estimated at >1 million generations for all species pairs (Figure 1C).

The Sespe Creek population (S25) is clearly a distinct lineage well outside of the *M. brevipes* trio. Although S25 is most closely related to *M. constrictus*, high genetic divergence between these lineages (*d_s_* = 0.06) suggest that S25 is a distinct species (Figure 1D, Table S4). Other species in this clade are not sampled and could be closer relatives to Sespe Creek, but none are known to occur in that geographic area. *M. nanus* is recovered as an outgroup to the other *Eunanus* section species as expected, with the *M. aurantiacus* complex set as the ultimate outgroup (Figure 1C). Species-level relationships from the maximum-likelihood tree were corroborated by a neighbor-joining tree and ASTRAL consensus tree (Figure S1).

Despite clear divergence between these *Mimulus* species, we also discovered substantial historical introgression. Using ABBA-BABA tests (Table 1, Table S5) and associated *F* statistics (Figures 2A and S2), TWISST (Figure S3), and TreeMix (Figure S4), we detected three signals of gene flow that were consistent across all methods. The first of these signals supports directional introgression from the *M. brevipes-M. fremontii-M. johnstonii* clade into the Sespe Creek population (Table 1: *D* = 0.135-0.149; Figure 2A: *F_branch_* = 4.8-5.9%, Figure S3: TWISST quartets 41.1/41.8/17.0, Figure S4: TreeMix m=0.193). A second, weaker signal of introgression occurs in the reciprocal direction from the *M. constrictus*-Sespe Creek clade into the *M. brevipes-M. fremontii-M. johnstonii* clade (Table 1: *D* = 0.027-0.046, Figure 2A: *F_branch_* = 0.4- 1.3%, Figure S3: TWISST quartets 70.3/15.7/14.0, Figure S4: TreeMix m=0.021). Finally, a third signal of introgression was detected between *M. nanus* and the clade containing *M. constrictus* and Sespe Creek (Table 1: *D* = 0.101-0.178, Figure 2A: *F_branch_* = 2.4-3.4%, Figure S3: TWISST quartets 73.5/15.2/11.3, Figure S4: TreeMix m=0.296). We summarize these patterns in Figure S5, but note that shared evolutionary history makes it challenging to assign introgression events to single lineages.

**Table 1.**
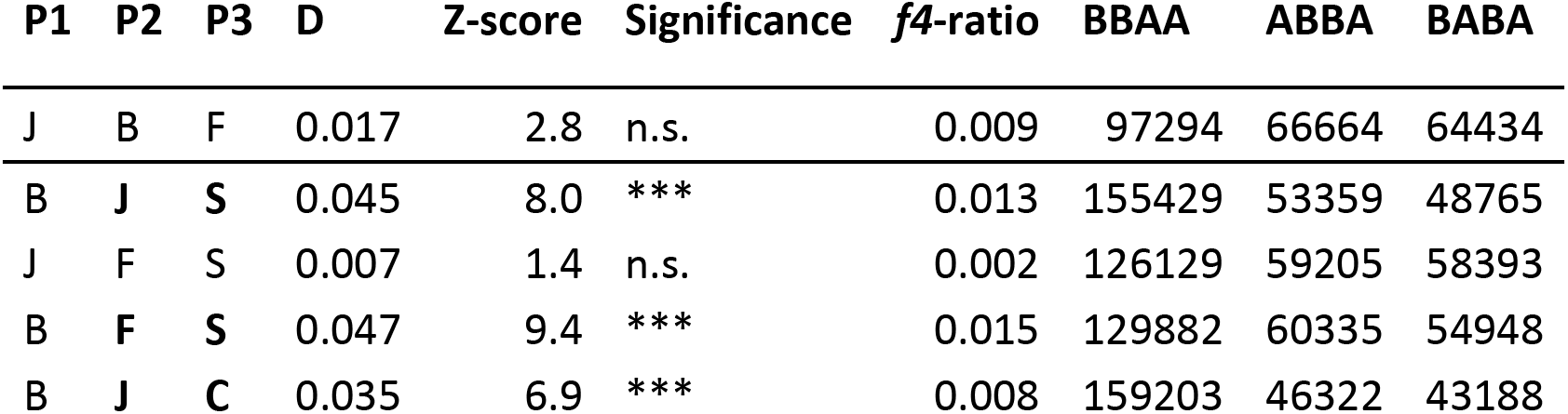
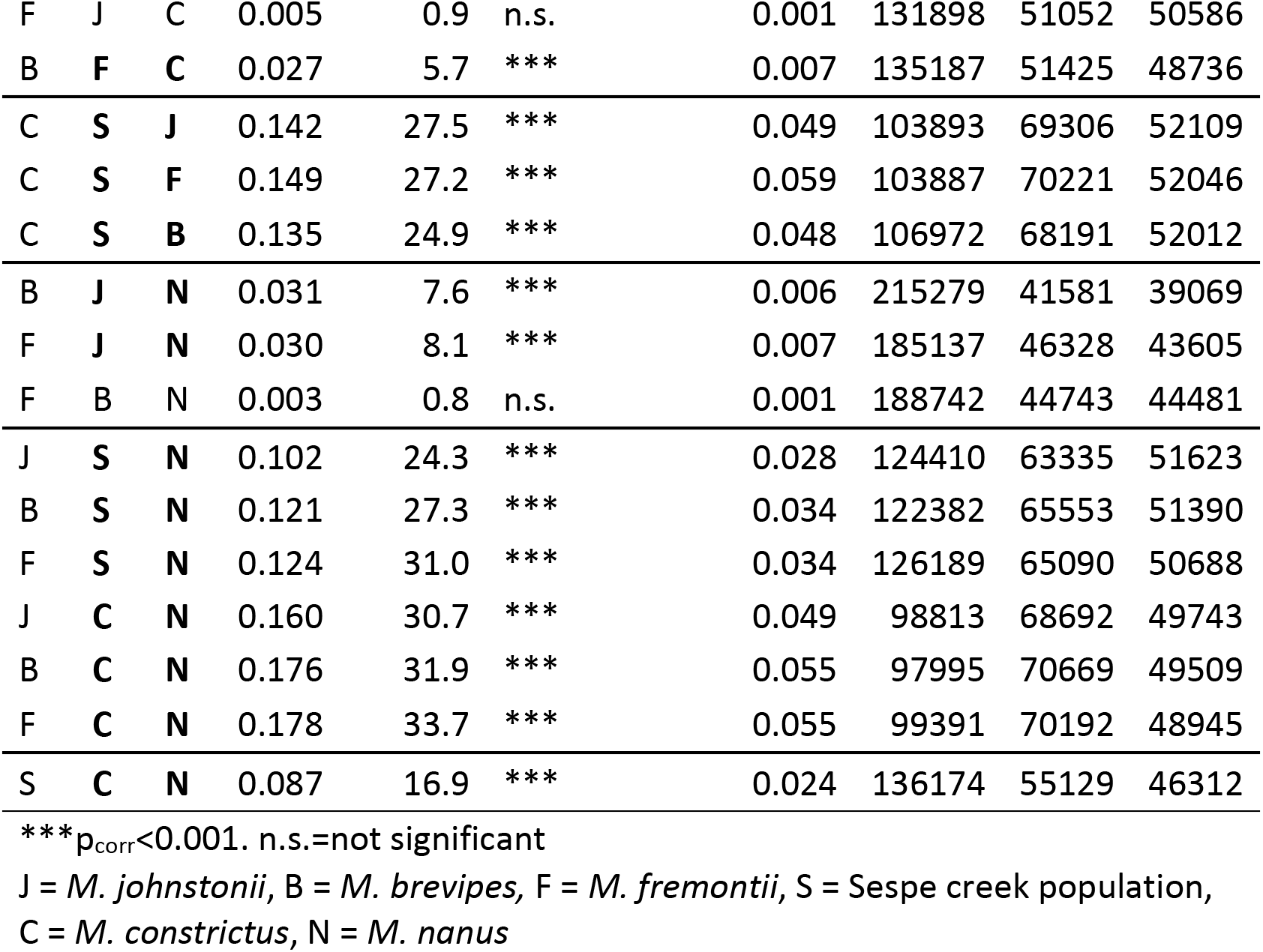
ABBA-BABA test results from downsampled SNP dataset.

| P1 | P2 | P3 | D | Z-score | Significance | f4-ratio | BBAA | ABBA | BABA |
| --- | --- | --- | --- | --- | --- | --- | --- | --- | --- |
| J | B | F | 0.017 | 2.8 | n.s. | 0.009 | 97294 | 66664 | 64434 |
| B | J | S | 0.045 | 8.0 | *** | 0.013 | 155429 | 53359 | 48765 |
| J | F | S | 0.007 | 1.4 | n.s. | 0.002 | 126129 | 59205 | 58393 |
| B | F | S | 0.047 | 9.4 | *** | 0.015 | 129882 | 60335 | 54948 |
| B | J | C | 0.035 | 6.9 | *** | 0.008 | 159203 | 46322 | 43188 |
| F | J | C | 0.005 | 0.9 | n.s. | 0.001 | 131898 | 51052 | 50586 |
| B | F | C | 0.027 | 5.7 | *** | 0.007 | 135187 | 51425 | 48736 |
| C | S | J | 0.142 | 27.5 | *** | 0.049 | 103893 | 69306 | 52109 |
| C | S | F | 0.149 | 27.2 | *** | 0.059 | 103887 | 70221 | 52046 |
| C | S | B | 0.135 | 24.9 | *** | 0.048 | 106972 | 68191 | 52012 |
| B | J | N | 0.031 | 7.6 | *** | 0.006 | 215279 | 41581 | 39069 |
| F | J | N | 0.030 | 8.1 | *** | 0.007 | 185137 | 46328 | 43605 |
| F | B | N | 0.003 | 0.8 | n.s. | 0.001 | 188742 | 44743 | 44481 |
| J | S | N | 0.102 | 24.3 | *** | 0.028 | 124410 | 63335 | 51623 |
| B | S | N | 0.121 | 27.3 | *** | 0.034 | 122382 | 65553 | 51390 |
| F | S | N | 0.124 | 31.0 | *** | 0.034 | 126189 | 65090 | 50688 |
| J | C | N | 0.160 | 30.7 | *** | 0.049 | 98813 | 68692 | 49743 |
| B | C | N | 0.176 | 31.9 | *** | 0.055 | 97995 | 70669 | 49509 |
| F | C | N | 0.178 | 33.7 | *** | 0.055 | 99391 | 70192 | 48945 |
| S | C | N | 0.087 | 16.9 | *** | 0.024 | 136174 | 55129 | 46312 |

\*\*\* $p_{\text{corr}} < 0.001$ . n.s.=not significant
J = *M. johnstonii*, B = *M. brevipes*, F = *M. fremontii*, S = Sespe creek population, C = *M. constrictus*, N = *M. nanus*

**Figure 2.**
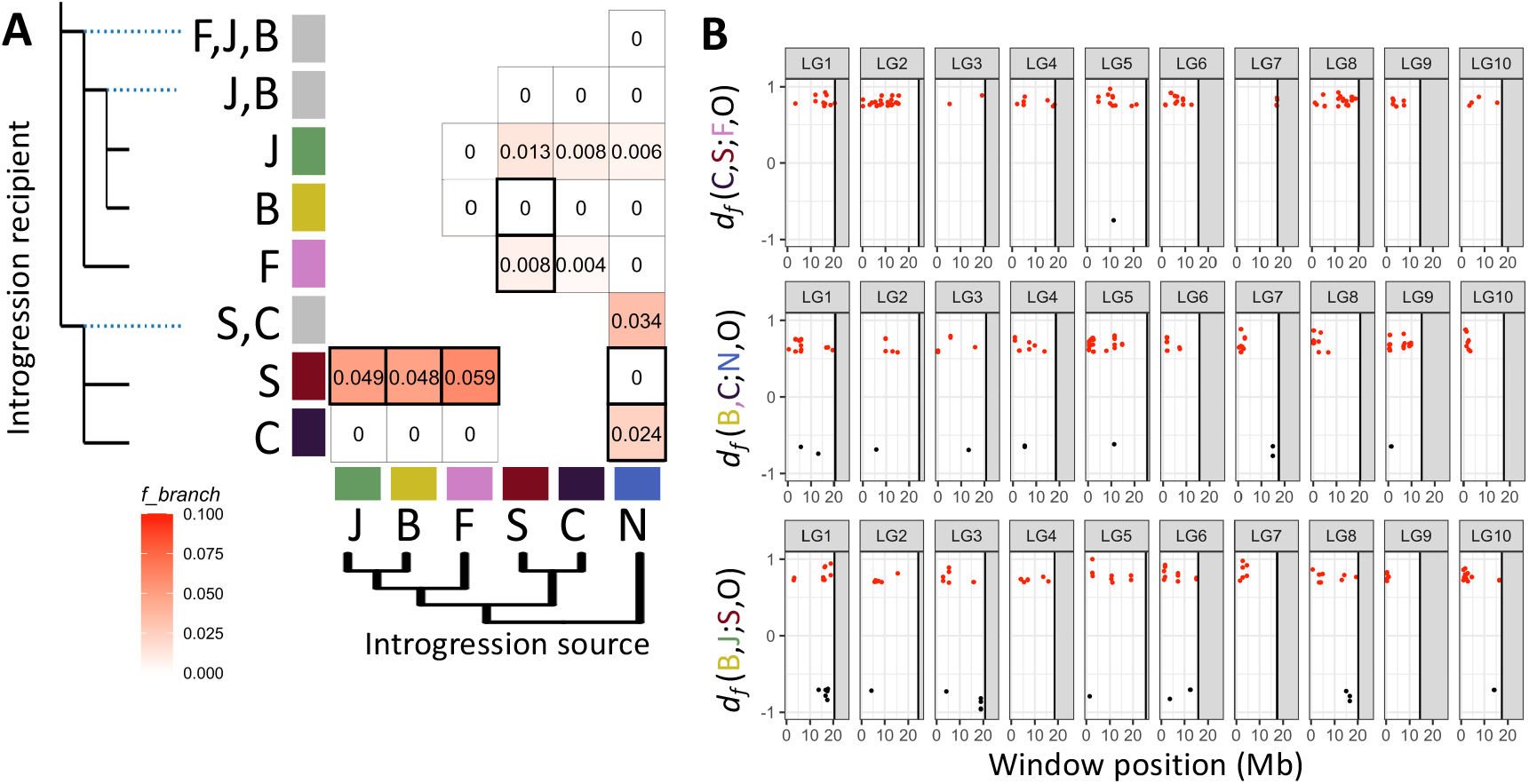
Signatures of historical introgression between species in *Mimulus* section *Eunanus*. **(A)** Genome-wide *F_branch_* statistics, an estimate of the proportion of the genome affected by introgression based on correlated *f4* statistics, using the downsampled SNP dataset. All listed nonzero *F_branch_* values are supported by ABBA-BABA tests with Bonferroni-corrected p-value < 0.001. Seven bold-outlined boxes indicate comparisons with significant gene discordance bias supporting introgression from TWISST gene tree summaries, using Bonferroni-corrected p- values <0.01. Full TWISST summaries are in Figure S3. (B) Position along ten linkage groups of extreme *df* values in 100-SNP windows for each of three tested quartets. Each quartet (P1,P2;P3,O) tests for signatures of introgression from P3 into P2 (positive outliers) or P3 into P1 (negative outliers). The top 100 windows with the largest absolute value of *df* are plotted, with positive *df* values in red and negative in black. From top to bottom, positive outliers show a history of introgression from *M. fremontii* into Sespe Creek, from *M. nanus* into *M. constrictus,* and from Sespe Creek into *M. johnstonii*. Total tested windows for each quartet: 9594, 10429, and 14596, respectively.

For each of these three general cases, we chose the trio with the strongest signal and scanned the genome for signatures of introgression using the statistic *df*. Outlier *df* windows were asymmetric, with more positive outliers (signals of introgression consistent with the genome-wide *F_branch_* pattern) than negative outliers of the same magnitude (Figure 2B). Positive outliers were widely distributed throughout the genome, present on all ten linkage groups for all three trios and with no large blocks of uninterrupted introgression, indicating that gene flow in this group is ancient rather than ongoing.

### Multiple strong postmating reproductive barriers between species in the *M. brevipes* group

Most crosses between four focal species (*M. johnstonii, M. brevipes, M. fremontii,* and the Sespe Creek population) showed little to no reduction in crossing success (probability of producing a seed, or seeds produced per fruit) compared to within-species crosses (Figures 3 and S6, Table S6). In one case (FxB), there is evidence of strong unidirectional postmating prezygotic isolation: only one in 58 crosses between maternal *M. fremontii* and paternal *M. brevipes* produced any seeds, a significantly lower proportion than the within-species or reciprocal cross types (Figure 3, Table S6). This pattern appears to be additive on both the maternal and paternal sides in *M. brevipes* x *M. fremontii* F1 hybrids, with intermediate crossing success in both HxB and FxH crosses (Figure S7). A few other interspecific cross types also had low or zero seed production (e.g., reciprocal crosses of *M. fremontii* and Sespe Creek), which could indicate a postmating prezygotic barrier, but these crosses had low sample sizes and did not reach significance compared to within-species crosses.

**Figure 3.**
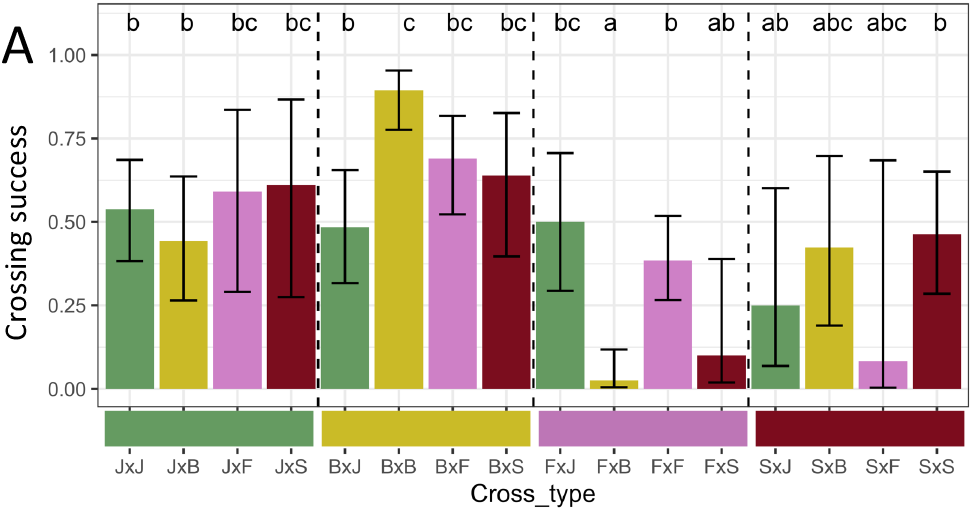
Postmating prezygotic isolation between at least one species pair. Crossing success (probability of producing at least one seed from a cross) for intra- and interspecific cross types. Shown are least-square means and asymptotic confidence intervals from a Firth binomial model. Letters indicate significance in post-hoc Tukey tests; cross types sharing a letter are not significantly different. One interspecific cross type, FxB, was significantly reduced compared to both intraspecific parental crosses, indicating an asymmetric barrier to fruit set. Two other crosses, FxS and SxF, had low fruit set possibly indicating a reproductive barrier, but were not significant in the model due to low sample sizes; e.g., 0 out of 4 SxF crosses resulted in seeds. J = *M. johnstonii*, B = *M. brevipes,* F = *M. fremontii*, S = Sespe creek population.

For crosses that did produce seeds, we found strong F1 hybrid seed inviability (viability <2%) in nine interspecific crosses (Figure 4A-B, Table S6). The exception was species pair *M. fremontii* and *M. brevipes,* which produced viable seeds in both directions (Figure 4A-B, Table S6). We could not score seed viability for cross type SxF because the only potential seeds produced were not distinguishable from unfertilized ovules.

**Figure 4.**
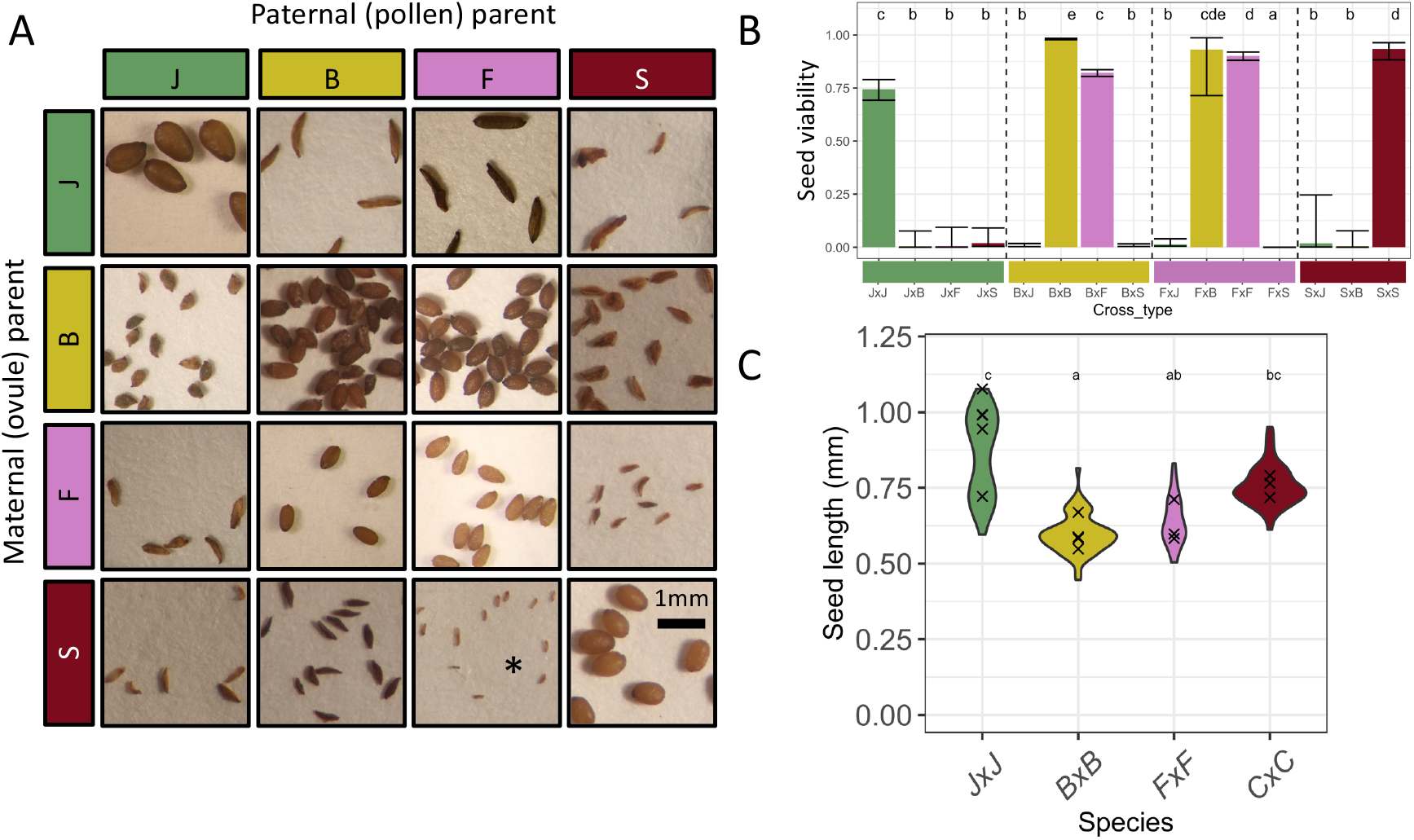
Strong hybrid seed inviability between multiple species pairs. (A) Representative seed images from intra- and interspecific laboratory crosses between four species. All images are adjusted to the same scale. For the SxF cross, marked with an asterisk, the only ‘seeds’ produced were not distinguishable from unfertilized ovules and were not counted as seeds for crossing success or viability scoring, but it is possible they represent early seed abortion. (B) Proportion of viable seeds produced by intra- and interspecific laboratory crosses. Shown are least-square means and asymptotic confidence intervals from a Firth binomial model, with letters indicating significance in post-hoc Tukey tests. Intraspecific, BxF, and FxB crosses are mostly viable, but all other interspecific cross types have almost complete inviability. (C) Seed lengths for intraspecific crosses. Violin plots show distribution of data for individual seeds, with x’s marking the means of individual fruits. Letters indicate significance from post-hoc Tukey tests of an LMM.

*M. johnstonii* intraspecific seeds were significantly larger than *M. brevipes* or *M. fremontii* seeds, with Sespe Creek seeds intermediate in size (Figure 4C, Table S7). Inviable seeds overall showed no evidence of overgrowth or undergrowth relative to parents, but instead tended to match the seed length of the maternal parent. However, F1 hybrids with Sespe Creek as maternal parent did have slightly smaller seeds than either parent (this trend was only significant for SxF) (Figures 4A and S8A, Table S7). Crosses with *M. johnstonii*, the largest-seeded species, as the maternal parent produced inviable seeds that were collapsed and thin, as shown by a significant increase in hybrid seed length/width ratios (three cross types) compared to both parents and to the reciprocal cross type (Figures 4A and S8B, Table S7). In the reciprocal direction, hybrid seeds were typically shriveled, as were hybrid seeds in both directions of Sespe Creek crosses (Figures 4A and S8B).

Although in most cases, strong seed inviability prevented us from quantifying later-acting barriers, we were able to assess F1 hybrid sterility between *M. brevipes* and *M. fremontii*. Pollen number per flower was intermediate for hybrids compared to the two parental species (Figure 5A, Table S8). However, only ∼20% of hybrid pollen grains were fertile, a sharp reduction from either parent, indicating strong F1 male sterility (Figure 5B, Table S8). We find no evidence of hybrid female sterility in *M. brevipes* x *M. fremontii* F1s; seed production from pollinated hybrids was intermediate compared to pure species (Figure 5C).

**Figure 5.**
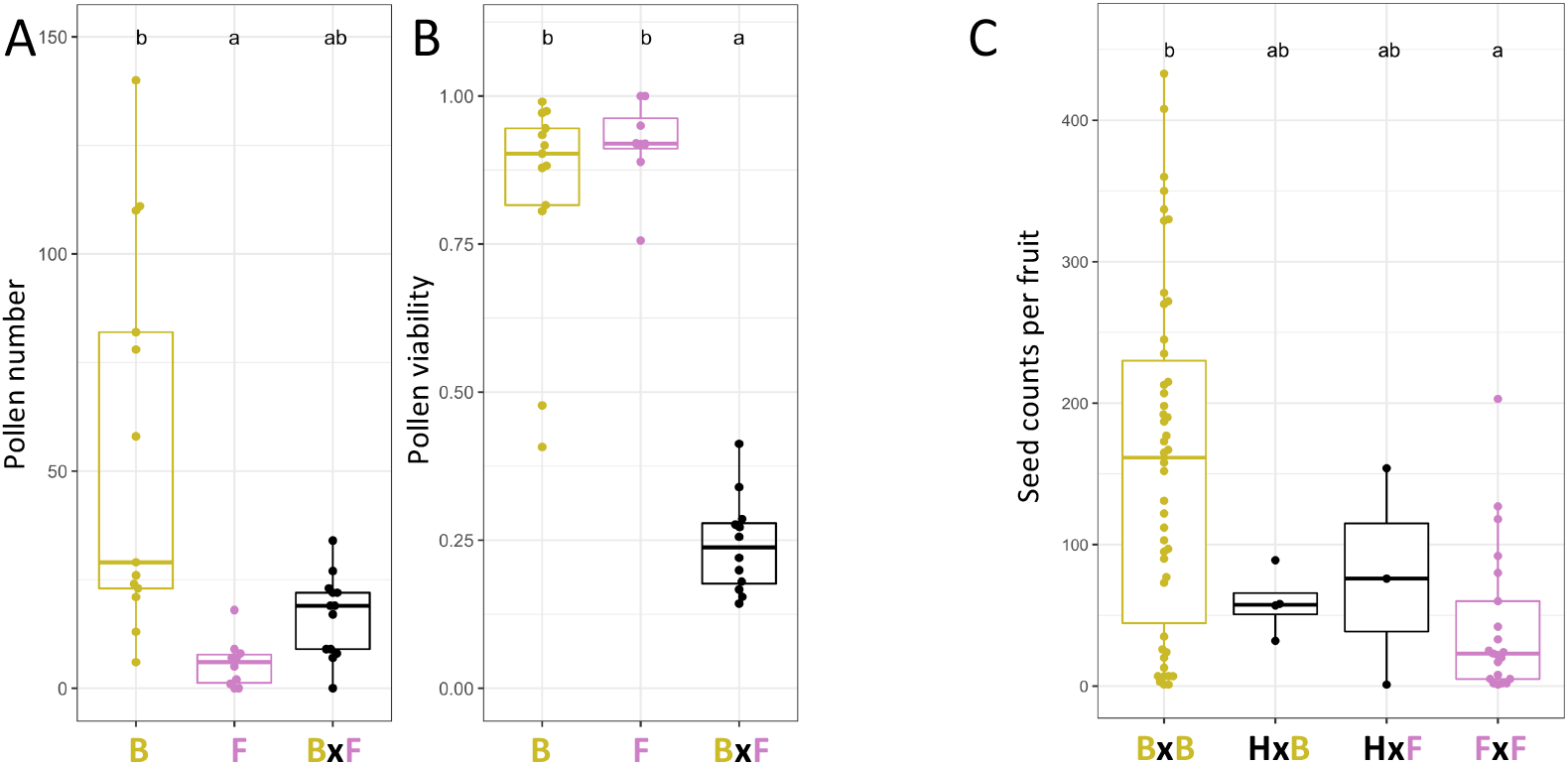
Strong hybrid male sterility in hybrids between *M. brevipes* and *M. fremontii*. (A) Pollen number (counts per mm^2^ on a hemocytometer) and (B) pollen viability (proportion of pollen grains scored as viable in aniline blue stain). Letters indicate significance in post-hoc Tukey tests from a poisson (abundance) or binomial (viability) GLMM with population as a random variable. Each point represents the average across 1-3 flowers from a single individual. For each panel, groups sharing a letter are not significantly different. (C) Seed production from hybrid vs. parental fruits as a test of hybrid female sterility, including only fruits that produced at least one seed. Letters indicate significance in post-hoc Tukey tests from a linear model. B=*M. brevipes*, F=*M. fremontii*, H=*M. brevipes x M. fremontii* F1 hybrid.

Total postmating reproductive isolation was near-complete (RI>=0.95) for all tested species pairs except for *M. brevipes* vs. *M. fremontii,* primarily driven by strong hybrid seed inviability (Table 2). The exception, *M. brevipes* vs. *M. fremontii,* had incomplete but still substantial total postmating RI from a combination of postmating prezygotic isolation and F1 male sterility (B > F: RI=0.91; F > B: RI=0.66).

**Table 2.** Summary of Postmating Reproductive Isolation (RI).

| Pair a, b | Crossing success RI |  | Seed viability RI |  | F1 viability RI |  | Total measured RI |  |
| --- | --- | --- | --- | --- | --- | --- | --- | --- |
|  | a>b | b>a | a>b | b>a | a>b | b>a | a>b | b>a |
| J, B | 0.30 | 0.10 | **0.99** | **1.00** | -- | -- | **0.99** | **1.00** |
| J, F | -0.13 | -0.05 | **0.97** | **1.00** | -- | -- | **0.97** | **1.00** |
| B, F | **0.91** | 0.13 | -0.03 | 0.09 | -- | **0.57** | **0.91** | **0.66** |
| J, S | 0.35 | -0.07 | **1.00** | **0.97** | -- | -- | **1.00** | **0.96** |
| B, S | 0.05 | 0.16 | **1.00** | **0.99** | -- | -- | **1.00** | **0.99** |
| F, S | 1.00 | 0.68 | -- | **1.00** | -- | -- | 1.00 | **1.00** |

a>b indicates reproductive isolation preventing gene flow from species a into species b, calculated by comparing the fitness of b x a crosses relative to b x b crosses, with b>a indicating the reciprocal direction using the fitness of a x b crosses relative to a x a crosses, following (Sobel & Chen, 2014).
**Bolded** values include contributions from barriers with significant model results.

## DISCUSSION

A major goal of speciation research is to understand similarities and differences in how reproductive isolation emerges as groups of species diverge. We find that *Mimulus brevipes, Mimulus johnstonii, and Mimulus fremontii* are clearly delineated species both genetically and by intrinsic reproductive barriers. We find strong hybrid seed inviability between multiple species pairs, as well as a postmating prezygotic barrier and severe hybrid male sterility between one species pair. While premating barriers have not yet been thoroughly quantified in this group, the combination of close geographic proximity (within centimeters in some cases) and similar floral morphology (excluding *M. brevipes*) makes it unlikely that premating barriers alone can eliminate hybridization. These strong postmating barriers are therefore likely to be important in preventing contemporary gene flow.

While many studies have shown that reproductive isolation increases with divergence on average, individual species pairs are often idiosyncratic (Coyne & Orr, 1989; Malone & Fontenot, 2008; Matute & Cooper, 2021; Moyle et al., 2004). Our results highlight this heterogeneity – while reproductive isolation and divergence are both strong overall, the species pair with the least postmating reproductive isolation in this group (*M. brevipes* and *M. fremontii*) is not the most recently diverged pair (*M. brevipes* and *M. johnstonii*). Paying greater attention to the exceptions may help us better understand the factors that drive reproductive isolation.

### Gene discordance and introgression

We find evidence of extensive phylogenetic discordance across the genome, including signatures of historical introgression between multiple species in this group. These signals are scattered throughout the genome and broken up by generations of recombination; we find no evidence of recent or ongoing introgression. Without further sampling of populations and species in this section, attributing these signals to precise events on the phylogeny is difficult. Taken as a whole, however, they demonstrate a high degree of complexity in the divergence process, consistent with reticulate evolutionary histories of other groups within *Mimulus* (e.g., Brandvain et al., 2014; Nelson et al., 2021; Stankowski et al., 2019), in plants more broadly (e.g., Curtu et al., 2009; Goulet et al., 2017; Hamlin et al., 2020; Kay et al., 2018; Scascitelli et al., 2010), and across the tree of life (e.g., D’Angiolo et al., 2020; Green et al., 2010; Kleindorfer et al., 2014; Mallet et al., 2007; Schumer et al., 2018). Without postmating reproductive barriers, even small amounts of introgression can lead to species collapse in sympatry, especially if premating barriers are disrupted by environmental change (Behm et al., 2010; Kleindorfer et al., 2014; Xiong & Mallet, 2022). Alternatively, species can persist despite ongoing introgression, depending on the circumstances (Kay et al., 2018; Kenney & Sweigart, 2016; Servedio & Hermisson, 2020). Past introgression in the *Eunanus* group may have predated the evolution of the strong postmating barriers we see today. We argue that postmating reproductive barriers likely played an important role in reducing gene flow as divergence increased, contradicting the hypothesis that postmating barriers are ‘after-the-fact’ byproducts with little influence on speciation trajectories.

### Cryptic diversity in Mimulus section Eunanus

Genetic data allow us to uncover cryptic variation in the form of distinct genetic lineages that are not readily distinguished by morphological features in the field (Bickford et al., 2007; Struck et al., 2018). Our population from Sespe Creek does not appear to match any of the likely species from the area, and we find that it is a genetically distinct lineage with a history of introgression. In addition, it appears to be reproductively isolated from other members of the group by strong hybrid seed inviability. We do not yet know the exact nature of its origin, or how widely distributed this lineage is; more sampling from the area and genetic data from more species in the group will be necessary to fully resolve these questions. Studies of speciation in plants often focus on ecologically mediated premating reproductive isolation, but cryptic diversity may be better explained by postmating barriers (Coughlan & Matute, 2020). *Mimulus* section *Eunanus* is home to many more understudied, small, pink-flowered taxa; if our study is any indication, strong postmating reproductive isolation may play an outsized role in generating diversity in this group.

### Parallel evolution of hybrid seed inviability in Mimulus

Hybrid seed inviability has been found repeatedly in species pairs from the *Mimulus guttatus* species complex (section *Simiolus*) (Coughlan et al., 2020; Oneal et al., 2016; Sandstedt & Sweigart, 2022). By demonstrating hybrid seed inviability in a distant section of *Mimulus,* we show that this pattern is widespread and important, both across *Mimulus* and likely in plants more broadly. In the *M. guttatus* complex and other systems, hybrid seed inviability has been tied to parental conflict in resource allocation, a process mediated by the endosperm (Coughlan et al., 2020; Haig & Westoby, 1991; Lafon-Placette et al., 2017; Oneal et al., 2016; Rebernig et al., 2015; Roth et al., 2019; Sandstedt & Sweigart, 2022). We do not find clear evidence of parental conflict in our system -- hybrid seed sizes tended to track maternal seed sizes, without the telltale overgrowth or undergrowth phenotypes. However, hybrid seed size is not always a good predictor of parental conflict (Sandstedt & Sweigart, 2022), so future work to characterize differences in seed development would be required to rule out a conflict hypothesis.

Alternatively, differences in parental seed sizes between our species could result in Dobhzansky-Muller-like incompatibilities during hybrid seed development, without invoking parental conflict as a driver. Seed size is an important life history trait tied to ecological strategies; larger seeds tend to have more success establishing under a variety of hazardous conditions, from nutrient deprivation to low soil moisture to deep burial (Leishman et al., 2000). Larger, heavier seeds may also prevent long-range dispersal and keep offspring in maternal habitats for which they are locally adapted, as is the case in dune-adapted plants (Bowers, 1982; Schwarzbach et al., 2001). Both *M. johnstonii* and the Sespe Creek population are found on unstable scree slopes, which may pose a particular challenge to seed establishment or favor local dispersal to maintain local adaptation. This habitat preference is therefore a possible candidate for a selective force, which could have acted in parallel in *M. johnstonii* and Sespe Creek to drive seed size differences leading to hybrid inviability.

### Postmating prezygotic isolation

Crossing failure (postmating prezygotic isolation) can be caused by a failure of pollen tube germination, failure of pollen tube growth, failure of fertilization, or even early seed abortion (Wheeler et al., 2001); it may be a passive incompatibility between pollen and pistil, or an active rejection mechanism to prevent maladaptive hybridization (Hogenboom et al., 1975; Roda & Hopkins, 2019; Rushworth et al., 2022). Most studies on the mechanisms of postmating prezygotic isolation come from systems with self-incompatibility, where they are thought to be related to self-incompatibility mechanisms, e.g. *Solanum* (Bernacchi & Tanksley, 1997; Tovar-Méndez et al., 2014), *Nicotiana* (Kuboyama et al., 1994), and *Lilium* (Ascher & Drewlow, 1975), though a mechanism unrelated to self-incompatibility has been described in *Brassica* (Fujii et al., 2019). Since *Mimulus* lacks self-incompatibility mechanisms, this group provides an opportunity to investigate how interspecific incompatibility may arise on its own. In other systems with differences in style length, germinating pollen tubes can under- or over-shoot the ovules (Gore et al., 1990; Williams & Rouse, 1988). Style length could be important in our system, since *M. brevipes* styles are substantially longer than *M. fremontii* styles, though our case is unusual in that pollen from the long-styled parent fails while pollen from the short-styled parent is successful. Pollen competition driving differential coevolution of the pollen and pistil is another possible source of incompatibilities (Brandvain & Haig, 2005; Skogsmyr & Lankinen, 2002), a scenario which has been documented in the *M. guttatus* complex (Aagaard et al., 2013; Fishman et al., 2008) and *Arabidopsis* (Takeuchi & Higashiyama, 2012).

### Hybrid male sterility

Hybrid male sterility has been characterized multiple times in *Mimulus* sections Simiolus and Erythranthe, with a variety of genetic causes, including a nuclear DMI interaction (Sweigart et al., 2006), a cytoplasmic-nuclear interaction (Fishman & Willis, 2006), or underdominant chromosomal rearrangements (Stathos & Fishman, 2014), but the relative frequency of these mechanisms across plants is unknown. *M. brevipes* and *M. fremontii* are more genetically divergent than most pairs for which male sterility has been studied, in *Mimulus* or elsewhere. Since they still produce viable hybrids and sterility is not 100% complete, genetic mapping could be used in the future to determine the mechanisms underlying sterility in this system, and the degree of parallelism with other cases in *Mimulus*.

### Future directions

As the first in-depth genomic and phenotypic investigation of speciation within *Mimulus* section *Eunanus,* this study fills an important phylogenetic gap within the speciation literature, helping to position the genus *Mimulus* as a leading model for broad phylogenetically informed comparisons of how species form and diverge. Our study system highlights both universal and idiosyncratic patterns in the speciation process; we add to a growing body of evidence that hybrid seed inviability and hybrid male sterility are important barriers to gene flow in *Mimulus* and across plant taxa; and we demonstrate the complexity of gene discordance and historical introgression among species despite substantial overall divergence. In addition, we lay the groundwork for fruitful avenues of future mechanistic study: seed size evolution and hybrid seed inviability; style length and postmating prezygotic isolation; and highly divergent hybrid male sterility are all worthy of future exploration in this group. Out study also provides the starting point for a more complete phylogenetic sampling of section *Eunanus*, which would enhance our understanding of cryptic divergence and the interplay between ecological, geographic, and genetic forces driving the diversification of species.

## Author contributions

Research conceived and designed jointly by M.C.F and A.L.S. Data collected and analyzed by M.C.F. Manuscript written by M.C.F and A.L.S.

## Data accessibility

All newly generated whole genome sequence data are deposited at the NCBI Sequence Read Archive (accession PRJNA922914). Other data will be made available on Dryad Digital Repository upon publication.

## Acknowledgments

We thank James Sobel for seed collections, for advice on germination and growth conditions, and for helpful discussions. Samuel Mantel, Makenzie Whitener, Natalie Gonzalez, and Emma Chandler provided comments that improved the manuscript. Kelly Dyer, Jill Anderson, Casey Bergman, Molly Schumer, John Willis, Jenn Coughlan, Elen Oneal, and Robert Franks also contributed to useful discussions. This work is supported by the National Science Foundation DEB1856180; the National Institutes of Health Predoctoral Training Grant 5T32GM007103, the Society for the Study of Evolution Lewontin Award, and a UGA Plant Center Doctoral Dissertation Improvement Grant.

The authors declare no competing interests.

## Supplementary Figures and Tables

**Figure S1.**
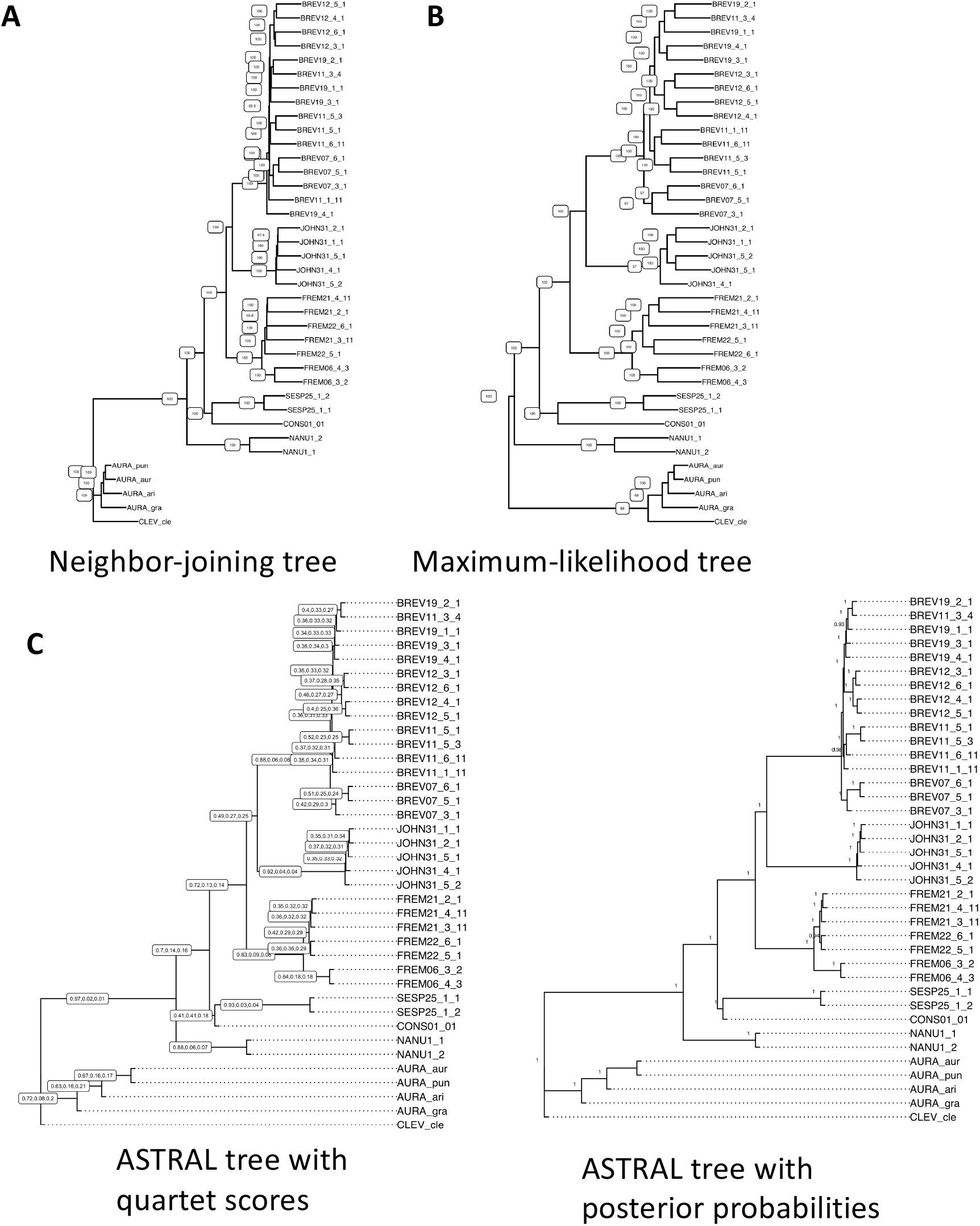
Comparison of phylogenetic methods. (A) Neighbor-joining tree produced from ‘complete’ SNP dataset, with heterozygous sites randomly assigned to one allele. (B) Maximum- likelihood tree produced using RAxML from the ‘complete’ SNP dataset using the GTR+Gamma model, with heterozygous sites randomly assigned to one allele. (C) ASTRAL tree produced from 17,524 gene trees, each created using RAxML with the GTR+Gamma model using SNPs from the ‘genic’ SNP dataset within a given gene’s coordinates, with heterozygous sites randomly assigned to one allele. Each node is labelled with the quartet support scores for the consensus, 1^st^ alternative, and 2^nd^ alternative topologies. (D) The same ASTRAL consensus tree shown in (C) but with posterior probabilities for each node.

**Figure S2.**
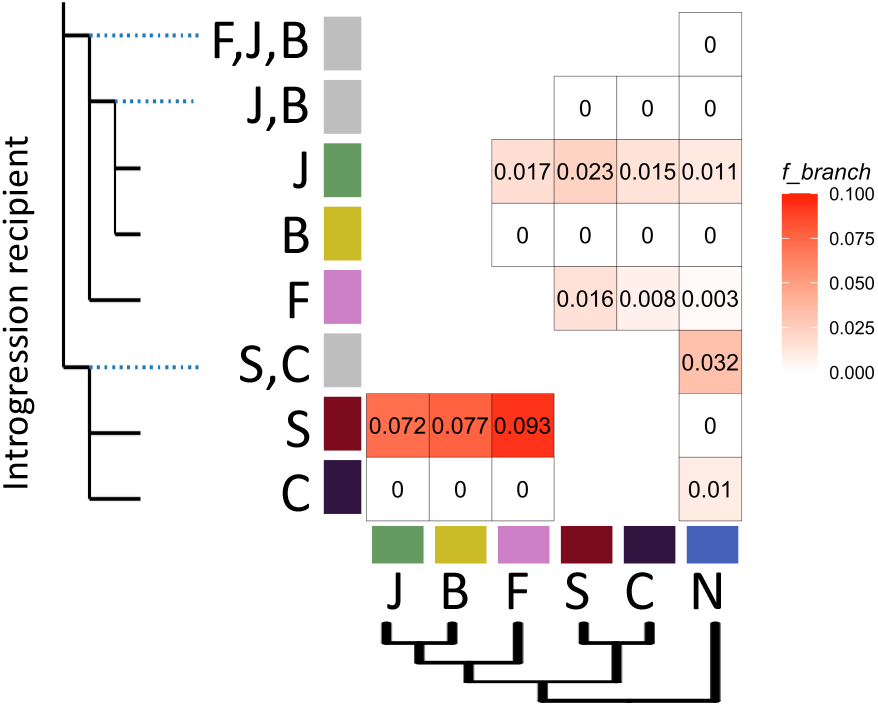
*F_branch_* results ‘complete’ SNP dataset. Results are qualitatively consistent with the ‘downsampled’ dataset (Fig 2) but estimated *F_branch_* values tend to be higher for the full dataset. J = *M. johnstonii*, B = *M. brevipes,* F = *M. fremontii*, S = Sespe creek population, C = *M. constrictus*, N = *M. nanus*.

**Figure S3.**
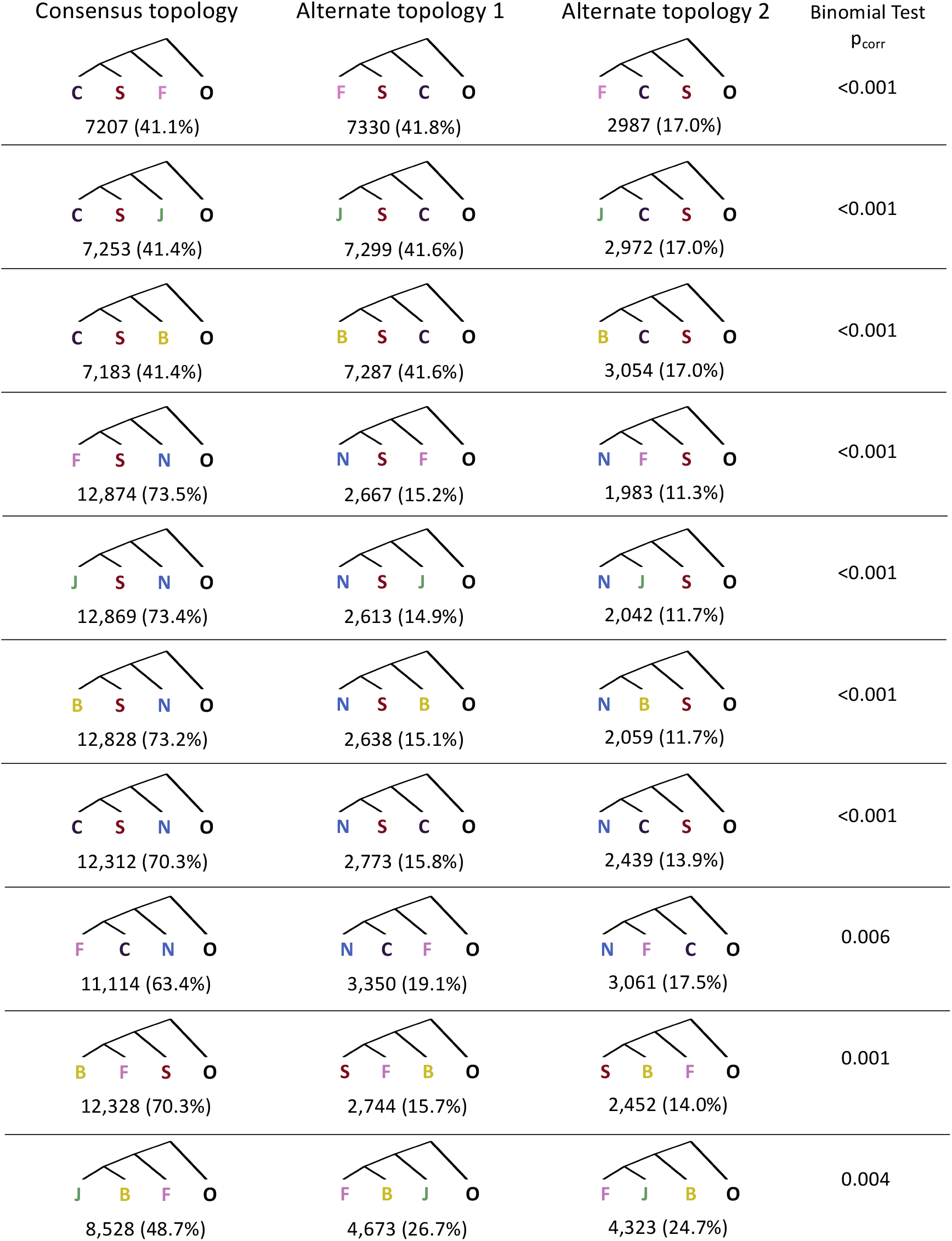
TWISST gene tree discordance summaries. Twenty trios were tested, using *M. aurantiacus* as the outgroup in all tests. Shown are the ten trios with significant overrepresentation of one alternate topology over another (binomial test, expected ratio 1:1, Bonferroni-corrected p_corr_<0.05). For each trio, the number of genes support each topology are given, along with the percentage of total trees. A total of 17,524 genes were used for all trios. J = *M. johnstonii*, B = *M. brevipes,* F = *M. fremontii*, S = Sespe creek population, C = *M. constrictus*, N = *M. nanus*, O = *M. aurantiacus* complex.

**Figure S4.**
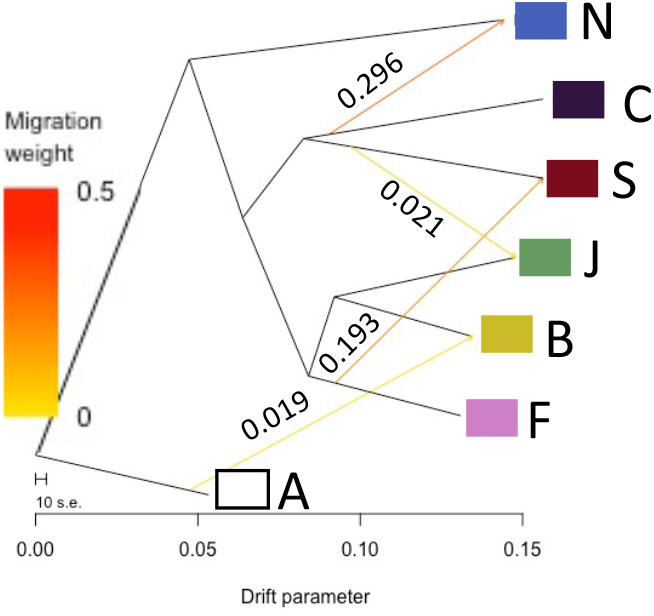
TreeMix network analysis results for four migration edges. The edge (C to N) was selected in the network with a single edge, which was the best model according to OptM; other edges improved the log-likelihood of the model by progressively decreasing amounts. Migration weights for the four-edge model are provided next to each edge.

**Figure S5.**
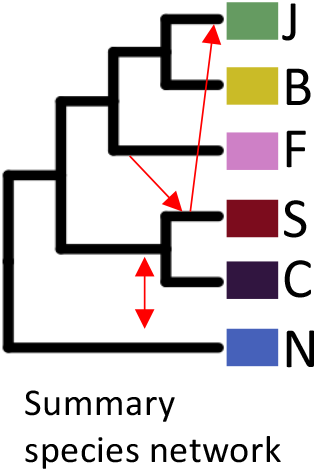
Summary of best-supported introgression events from multiple methods. Three introgression events were supported by *f_branch_ ,* TWISST, and TreeMix results. The exact identity and direction of individual introgression events may be different due to shared ancestry and incomplete sampling; for example, signatures of introgression from all of *M. fremontii, M. johnstonii, and M. brevipes* into Sespe Creek may be caused by a single event in a common ancestor, an unsampled relative, or just one of the sampled species.

**Figure S6.**
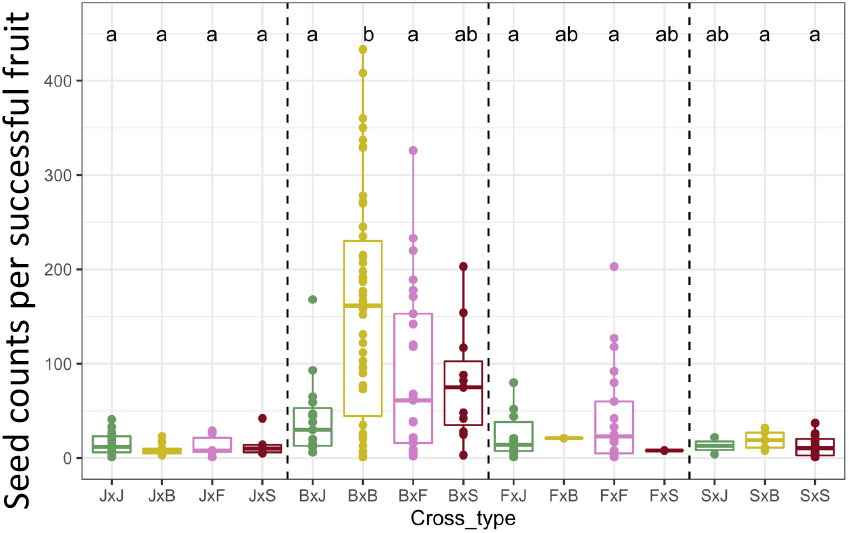
Seed counts do not indicate hybrid deficiency. Seed counts per fruit for fruits producing at least one seed. *M. brevipes* produces more seeds per fruit than the other species, but interspecific crosses do not show a reduction in seed count relative to intraspecific crosses. Letters indicate significance in post-hoc Tukey tests from a linear model; cross types sharing a letter are not significantly different.

**Figure S7.**
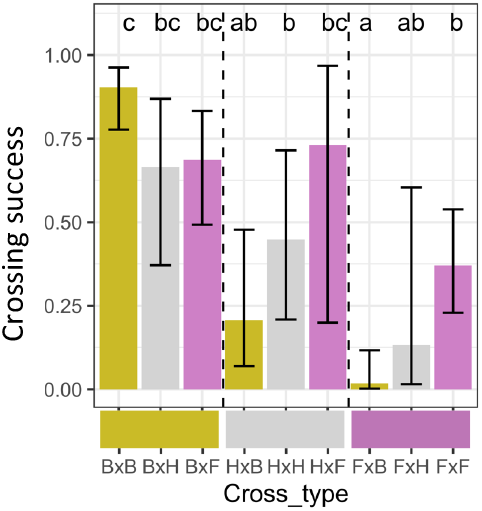
Crossing success in BxF hybrids is additive on maternal and paternal sides. Crossing success (probability of at least one seed produced from a cross) from crosses involving *M. brevipes* x *M. fremontii* F1 hybrids and their parental species. Shown are least-square means and asymptotic confidence intervals from a binomial GLMM. Letters indicate significance in post-hoc Tukey tests; cross types sharing a letter are not significantly different. HxB crosses have intermediate crossing success compared to BxB and FxB crosses (gold bars), indicating an additive effect of maternal species. Similarly, FxH crosses have intermediate crossing success compared to FxB and FxF crosses (right three bars), indicating an additive effect of paternal species. HxH have intermediate success compared to BxH and FxH (gray bars), as well as compared to HxB and HxF (middle three bars), further supporting additive patterns. B = *M. brevipes,* F = *M. fremontii*, H = F1 hybrids between *M. brevipes* and *M. fremontii*.

**Figure S8.**
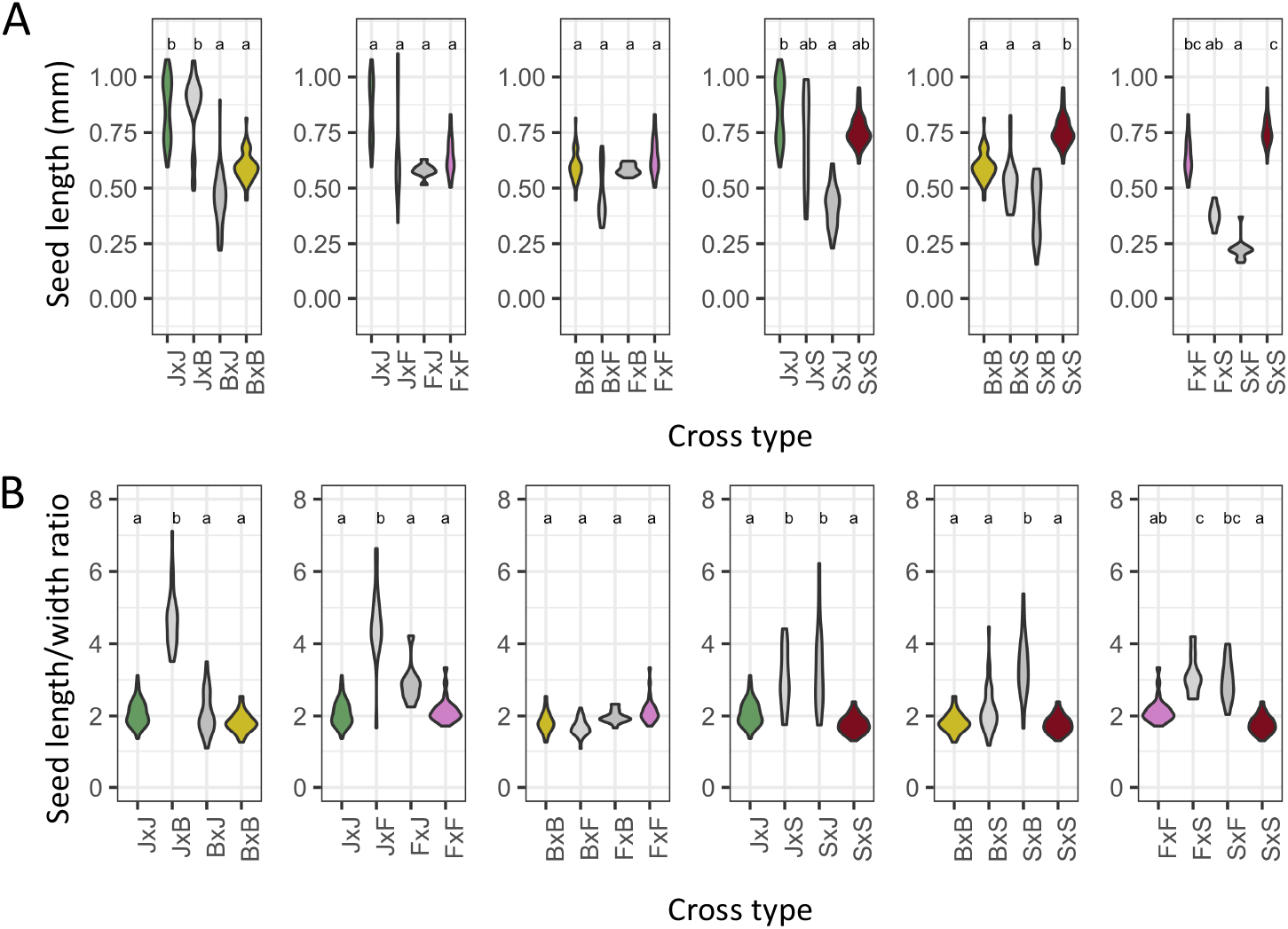
Seed measurements for all intra- and interspecific cross types. (A) Seed length and (B) length/width ratios. Violin plots show distribution of individual seeds. Letters indicate significance from post-hoc Tukey tests after running six independent species-pair models. One species pair had a significant reciprocal difference in hybrid seed size, while three pairs had a significant reciprocal difference in seed length/width ratio; length/width ratio was elevated compared to both parents for six cross types.

**Table S1.**
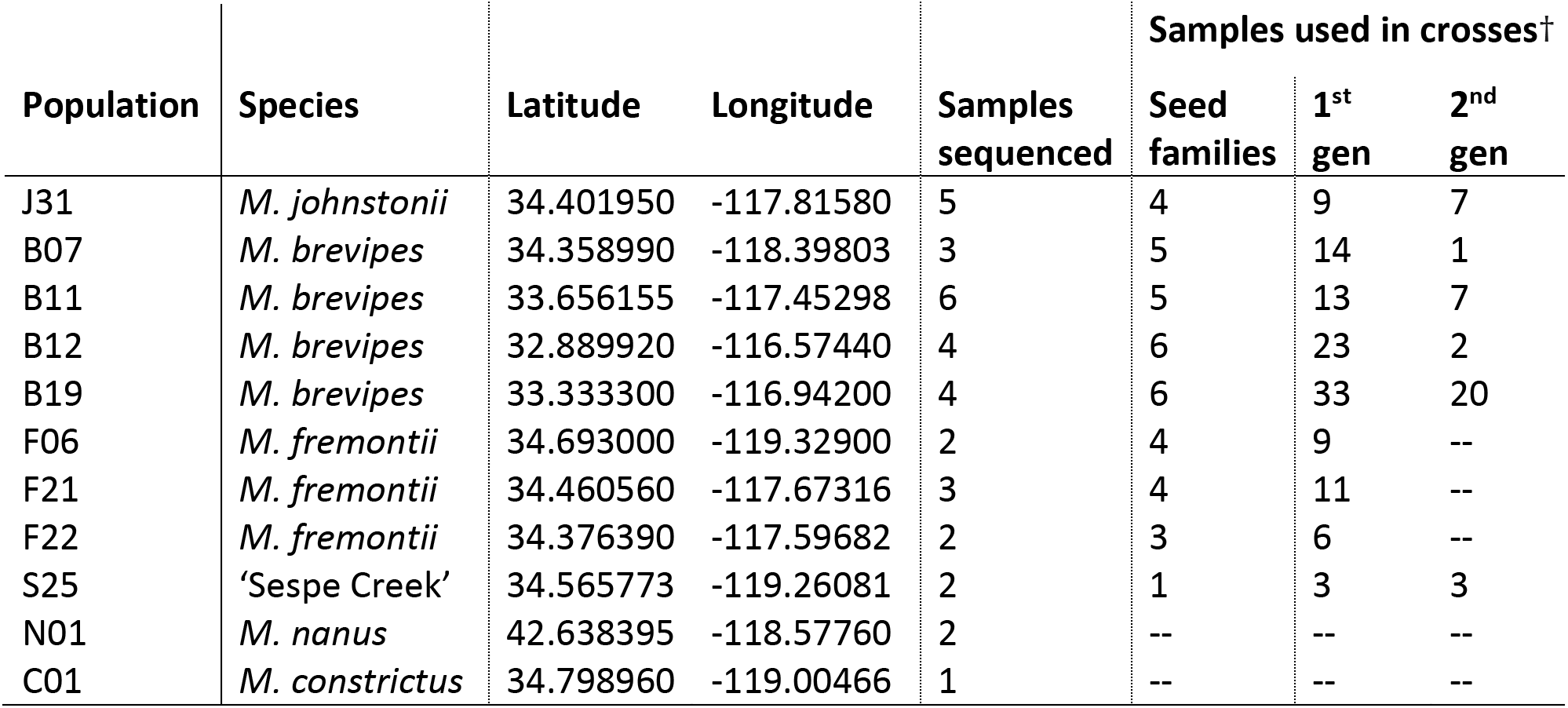
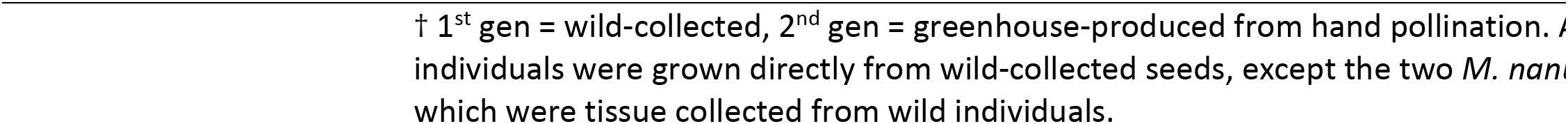
Population location information and number of samples used per population.

| Population | Species | Latitude | Longitude | Samples sequenced | Samples used in crosses† |  |  |
| --- | --- | --- | --- | --- | --- | --- | --- |
|  |  |  |  |  | Seed families | 1<sup>st</sup> gen | 2<sup>nd</sup> gen |
| J31 | *M. johnstonii* | 34.401950 | -117.81580 | 5 | 4 | 9 | 7 |
| B07 | *M. brevipes* | 34.358990 | -118.39803 | 3 | 5 | 14 | 1 |
| B11 | *M. brevipes* | 33.656155 | -117.45298 | 6 | 5 | 13 | 7 |
| B12 | *M. brevipes* | 32.889920 | -116.57440 | 4 | 6 | 23 | 2 |
| B19 | *M. brevipes* | 33.333300 | -116.94200 | 4 | 6 | 33 | 20 |
| F06 | *M. fremontii* | 34.693000 | -119.32900 | 2 | 4 | 9 | -- |
| F21 | *M. fremontii* | 34.460560 | -117.67316 | 3 | 4 | 11 | -- |
| F22 | *M. fremontii* | 34.376390 | -117.59682 | 2 | 3 | 6 | -- |
| S25 | ‘Sespe Creek’ | 34.565773 | -119.26081 | 2 | 1 | 3 | 3 |
| N01 | *M. nanus* | 42.638395 | -118.57760 | 2 | -- | -- | -- |
| C01 | *M. constrictus* | 34.798960 | -119.00466 | 1 | -- | -- | -- |

† 1<sup>st</sup> gen = wild-collected, 2<sup>nd</sup> gen = greenhouse-produced from hand pollination. All individuals were grown directly from wild-collected seeds, except the two *M. nanus* which were tissue collected from wild individuals.

**Table S2.** Sequencing and coverage information.

| Sample info |  |  | Read pairs |  |  | Coverage per base |  |
| --- | --- | --- | --- | --- | --- | --- | --- |
| Sample† | Pop | Batch | Raw | Aligned | % | All sites | % with >=4X coverage |
| BREV07_3_1 | B07 | 1 | 47,522,461 | 13,277,523 | 27.9% | 12.47 | 39.03% |
| BREV07_5_1 | B07 | 1 | 49,800,600 | 15,174,402 | 30.5% | 14.04 | 39.09% |
| BREV12_3_1 | B12 | 1 | 49,704,577 | 15,840,128 | 31.9% | 15.68 | 37.55% |
| BREV12_5_1 | B12 | 1 | 49,785,466 | 14,465,999 | 29.1% | 13.52 | 39.79% |
| BREV12_6_1 | B19 | 1 | 49,555,797 | 15,518,193 | 31.3% | 14.54 | 39.49% |
| BREV19_1_1 | B19 | 1 | 49,652,489 | 16,050,274 | 32.3% | 15.62 | 37.83% |
| FREM22_6_1 | F22 | 1 | 49,332,251 | 8,451,160 | 17.1% | 7.30 | 33.81% |
| JOHN31_5_1 | J31 | 1 | 49,714,033 | 13,727,590 | 27.6% | 12.82 | 37.51% |
| BREV07_6_1 | B07 | 2 | 25,682,314 | 5,672,576 | 22.1% | 5.67 | 32.30% |
| BREV11_5_3 | B11 | 2 | 28,911,480 | 5,804,484 | 20.1% | 5.35 | 31.82% |
| BREV12_4_1 | B12 | 2 | 28,052,293 | 6,443,496 | 23.0% | 6.28 | 33.38% |
| BREV19_2_1 | B19 | 2 | 29,339,237 | 7,002,395 | 23.9% | 6.87 | 33.83% |
| BREV19_3_1 | B19 | 2 | 36,190,791 | 3,248,019 | 9.0% | 3.10 | 21.10% |
| BREV19_4_1 | B19 | 2 | 14,858,533 | 2,429,889 | 16.4% | 2.19 | 16.18% |
| FREM06_3_2 | F06 | 2 | 30,075,452 | 5,408,560 | 18.0% | 4.51 | 26.45% |
| FREM06_4_3 | F06 | 2 | 29,784,495 | 6,422,159 | 21.6% | 5.68 | 29.74% |
| FREM21_2_1 | F21 | 2 | 33,601,408 | 5,451,858 | 16.2% | 4.60 | 27.10% |
| FREM22_5_1 | F22 | 2 | 24,172,306 | 2,679,095 | 11.1% | 2.35 | 19.37% |
| JOHN31_1_1 | J31 | 2 | 47,347,007 | 6,441,313 | 13.6% | 6.20 | 31.22% |
| JOHN31_2_1 | J31 | 2 | 64,312,829 | 10,195,170 | 15.9% | 9.88 | 36.43% |
| JOHN31_4_1 | J31 | 2 | 23,147,570 | 3,938,270 | 17.0% | 3.82 | 24.04% |
| JOHN31_5_2 | J31 | 2 | 32,146,268 | 5,382,624 | 16.7% | 4.86 | 25.48% |
| BREV11_1_11 | B11 | 3 | 12,194,294 | 3,311,065 | 27.2% | 3.16 | 18.64% |
| BREV11_3_4 | B11 | 3 | 16,149,111 | 4,165,590 | 25.8% | 3.88 | 24.46% |
| BREV11_5_1 | B11 | 3 | 19,609,920 | 5,018,757 | 25.6% | 4.72 | 26.70% |
| BREV11_6_11 | B11 | 3 | 17,240,947 | 4,349,611 | 25.2% | 4.02 | 26.63% |
| FREM21_3_11 | F21 | 3 | 31,307,714 | 5,636,788 | 18.0% | 4.86 | 30.25% |
| FREM21_4_11 | F21 | 3 | 23,442,361 | 4,186,587 | 17.9% | 3.67 | 24.90% |
| SESP25_1_1 | S25 | 3 | 37,881,366 | 8,233,912 | 21.7% | 7.70 | 34.70% |
| SESP25_1_2 | S25 | 3 | 22,013,442 | 5,056,136 | 23.0% | 4.58 | 29.93% |
| NANU01_1 | N01 | 3 | 15,545,612 | 2,612,978 | 16.8% | 2.26 | 14.32% |
| NANU01_2 | N01 | 3 | 14,729,812 | 2,603,428 | 17.7% | 2.38 | 17.61% |
| CONS01_1_1 | C01 | 4 | 16,909,453 | 4,072,713 | 24.1% | 4.10 | 16.6% |
| AURA_ari | AURA | SRX6077155 | 15,314,328 | 5,630,656 | 36.8% | 7.14 | 64.06% |
| AURA_aur | AURA | SRX6077153 | 18,453,062 | 7,584,963 | 41.1% | 10.02 | 75.92% |
| AURA_gra | AURA | SRX6077242 | 12,454,896 | 6,061,103 | 48.7% | 7.84 | 66.98% |
| AURA_pun | AURA | SRX6077404 | 14,056,153 | 5,076,335 | 36.1% | 6.53 | 60.90% |
| CLEV_cle | AURA | SRX6077240 | 12,810,520 | 4,567,552 | 35.7% | 5.58 | 53.01% |

† 1st number indicates population, 2nd number indicates maternal family, 3rd number refers to the individual plant. *M. nanus* samples (NANU01\_1 and NANU01\_2) are from distinct wild plants.

**Table S3.** Pedigree information for M. brevipes and M. fremontii F1 families.

| Line name | Line type | Maternal | Paternal | Number of individuals used |  |  |
| --- | --- | --- | --- | --- | --- | --- |
|  |  |  |  | Pollen viability | Maternal crosses | Paternal crosses |
| BF1 | BxF F1 hybrid | B19_3_2 | F6_4_3 | 0 | 1 | 1 |
| BF2 | BxF F1 hybrid | B12_5_2 | F21_3_1 | 2 | 1 | 1 |
| BF3 | BxF F1 hybrid | B19_1_1 | F6_3_1 | 7 | 7 | 6 |
| BF5 | BxF F1 hybrid | B19_4_14 | F21_4_11 | 4 | 2 | 0 |

**Table S4.** Nucleotide diversity and divergence values at synonymous sites. Values were calculated using pixy for all individual pairwise comparisons, then averaged across comparisons within a group.

| Species | Heterozygosity | Diversity |
| --- | --- | --- |
| AUR | 0.0025 | 0.0163 |
| B | 0.0198 | 0.0229 |
| C | 0.0135 | -- |
| F | 0.0209 | 0.0316 |
| J | 0.0170 | 0.0174 |
| N | 0.0154 | 0.0324 |
| S | 0.0180 | 0.0178 |
| Species 1 | Species 2 | Divergence |
| AUR | C | 0.0940 |
| AUR | N | 0.0945 |
| AUR | S | 0.0991 |
| AUR | B | 0.1000 |
| AUR | J | 0.1005 |
| AUR | F | 0.1027 |
| B | J | 0.0525 |
| B | F | 0.0599 |
| B | S | 0.0697 |
| B | C | 0.0707 |
| B | N | 0.0830 |
| C | S | 0.0605 |
| C | J | 0.0702 |
| C | F | 0.0730 |
| C | N | 0.0753 |
| F | J | 0.0598 |
| F | S | 0.0715 |
| F | N | 0.0860 |
| J | S | 0.0692 |
| J | N | 0.0828 |
| N | S | 0.0791 |

**Table S5.**
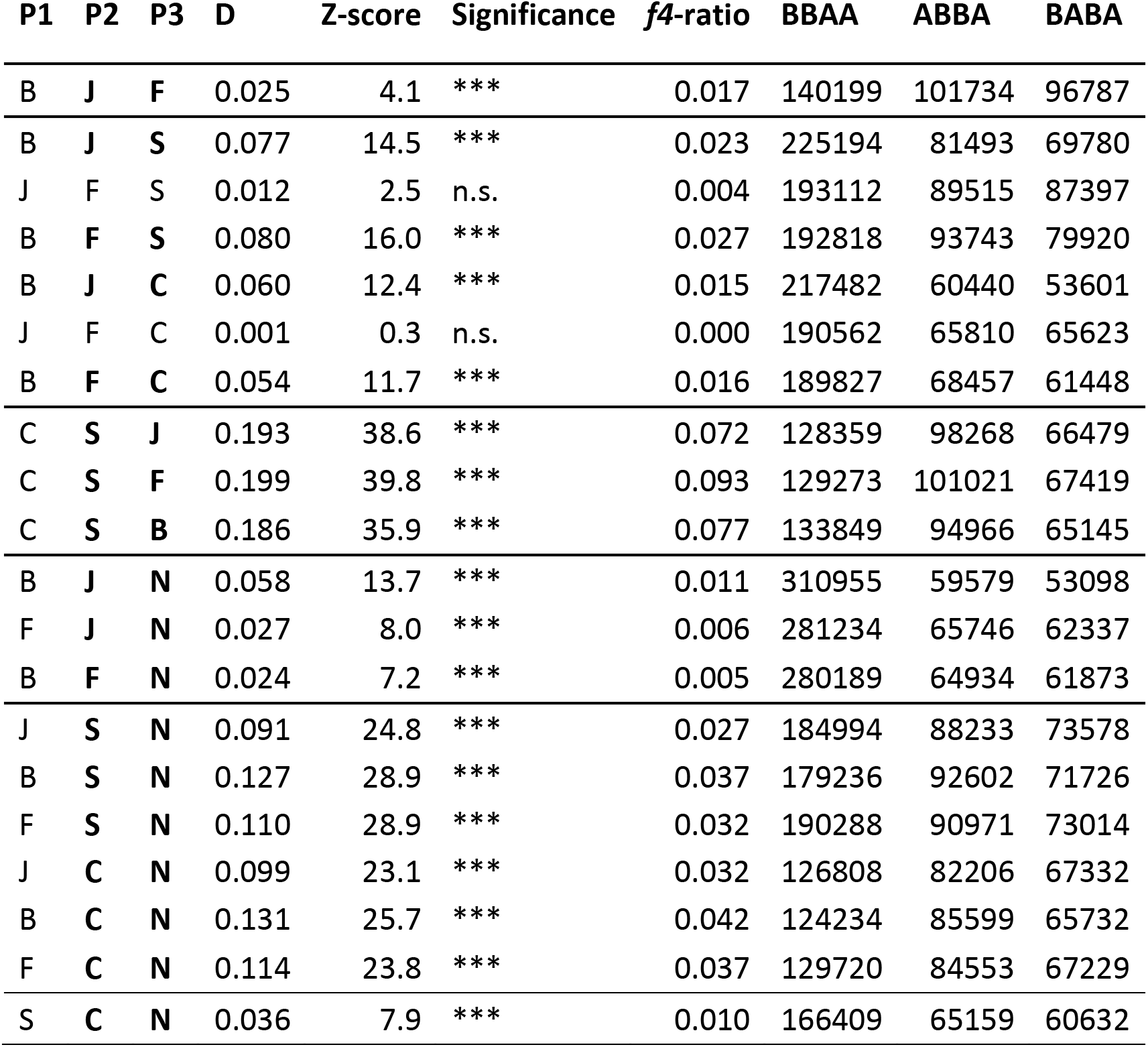
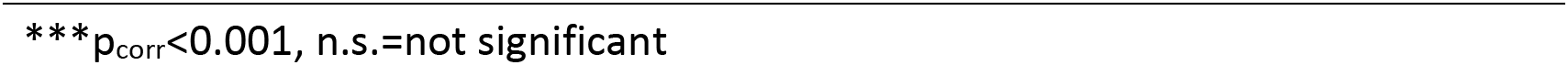
ABBA-BABA test results for the ‘complete’ SNP dataset.

| P1 | P2 | P3 | D | Z-score | Significance | *f*4-ratio | BBAA | ABBA | BABA |
| --- | --- | --- | --- | --- | --- | --- | --- | --- | --- |
| B | J | F | 0.025 | 4.1 | *** | 0.017 | 140199 | 101734 | 96787 |
| B | J | S | 0.077 | 14.5 | *** | 0.023 | 225194 | 81493 | 69780 |
| J | F | S | 0.012 | 2.5 | n.s. | 0.004 | 193112 | 89515 | 87397 |
| B | F | S | 0.080 | 16.0 | *** | 0.027 | 192818 | 93743 | 79920 |
| B | J | C | 0.060 | 12.4 | *** | 0.015 | 217482 | 60440 | 53601 |
| J | F | C | 0.001 | 0.3 | n.s. | 0.000 | 190562 | 65810 | 65623 |
| B | F | C | 0.054 | 11.7 | *** | 0.016 | 189827 | 68457 | 61448 |
| C | S | J | 0.193 | 38.6 | *** | 0.072 | 128359 | 98268 | 66479 |
| C | S | F | 0.199 | 39.8 | *** | 0.093 | 129273 | 101021 | 67419 |
| C | S | B | 0.186 | 35.9 | *** | 0.077 | 133849 | 94966 | 65145 |
| B | J | N | 0.058 | 13.7 | *** | 0.011 | 310955 | 59579 | 53098 |
| F | J | N | 0.027 | 8.0 | *** | 0.006 | 281234 | 65746 | 62337 |
| B | F | N | 0.024 | 7.2 | *** | 0.005 | 280189 | 64934 | 61873 |
| J | S | N | 0.091 | 24.8 | *** | 0.027 | 184994 | 88233 | 73578 |
| B | S | N | 0.127 | 28.9 | *** | 0.037 | 179236 | 92602 | 71726 |
| F | S | N | 0.110 | 28.9 | *** | 0.032 | 190288 | 90971 | 73014 |
| J | C | N | 0.099 | 23.1 | *** | 0.032 | 126808 | 82206 | 67332 |
| B | C | N | 0.131 | 25.7 | *** | 0.042 | 124234 | 85599 | 65732 |
| F | C | N | 0.114 | 23.8 | *** | 0.037 | 129720 | 84553 | 67229 |
| S | C | N | 0.036 | 7.9 | *** | 0.010 | 166409 | 65159 | 60632 |

\*\*\*p<sub>corr</sub><0.001, n.s.=not significant

**Table S6.** Crossing success and seed viability for each cross type.

| Maternal species | Paternal species | Cross type | Crossing success (total crosses) | Seed viability (total seeds) |
| --- | --- | --- | --- | --- |
| *M. johnstonii* | *M. johnstonii* | JxJ | 0.538 (39) | 0.745 (306) |
| *M. johnstonii* | *M. brevipes* | JxB | 0.440 (25) | **0.000 (98)** |
| *M. johnstonii* | *M. fremontii* | JxF | 0.600 (10) | **0.000 (78)** |
| *M. johnstonii* | Sespe Creek | JxS | 0.600 (5) | **0.015 (66)** |
| *M. brevipes* | *M. johnstonii* | BxJ | 0.484 (31) | **0.006 (630)** |
| *M. brevipes* | *M. brevipes* | BxB | 0.902 (51) | 0.981 (7292) |
| *M. brevipes* | *M. fremontii* | BxF | 0.694 (36) | 0.821 (2226) |
| *M. brevipes* | Sespe Creek | BxS | 0.500 (6) | **0.018 (217)** |
| *M. fremontii* | *M. johnstonii* | FxJ | 0.500 (20) | **0.012 (246)** |
| *M. fremontii* | *M. brevipes* | FxB | **0.017 (58)** | 0.952 (21) |
| *M. fremontii* | *M. fremontii* | FxF | 0.382 (55) | 0.902 (911) |
| *M. fremontii* | Sespe Creek | FxS | 0.091 (11) | **0.000 (8)** |
| Sespe Creek | *M. johnstonii* | SxJ | 0.125 (8) | **0.000 (4)** |
| Sespe Creek | *M. brevipes* | SxB | 0.333 (6) | **0.000 (38)** |
| Sespe Creek | *M. fremontii* | SxF | 0.000 (4) | n/a (0) |
| Sespe Creek | Sespe Creek | SxS | 0.409 (22) | 0.909 (88) |

**Bolded** values are significant in Tukey tests compared to both parental cross types.

**Table S7.** Average seed measurements and sample sizes by cross type.

| Maternal species | Paternal species | Cross type | Seeds measured (number of fruits) | Average seed length (mm) | Average seed length/width ratio |
| --- | --- | --- | --- | --- | --- |
| *M. johnstonii* | *M. johnstonii* | JxJ | 67 (7) | 0.858 | 2.070 |
| *M. johnstonii* | *M. brevipes* | JxB | 50 (7) | 0.865 | 4.729 |
| *M. johnstonii* | *M. fremontii* | JxF | 26 (3) | 0.687 | 4.580 |
| *M. johnstonii* | Sespe Creek | JxS | 12 (1) | 0.726 | 3.038 |
| *M. brevipes* | *M. johnstonii* | BxJ | 66 (5) | 0.447 | 2.060 |
| *M. brevipes* | *M. brevipes* | BxB | 60 (4) | 0.597 | 1.813 |
| *M. brevipes* | *M. fremontii* | BxF | 45 (3) | 0.470 | 1.677 |
| *M. brevipes* | Sespe Creek | BxS | 56 (2) | 0.523 | 2.155 |
| *M. fremontii* | *M. johnstonii* | FxJ | 9 (2) | 0.579 | 2.901 |
| *M. fremontii* | *M. brevipes* | FxB | 15 (1) | 0.584 | 1.989 |
| *M. fremontii* | *M. fremontii* | FxF | 27 (3) | 0.641 | 2.149 |
| *M. fremontii* | Sespe Creek | FxS | 8 (1) | 0.377 | 3.134 |
| Sespe Creek | *M. johnstonii* | SxJ | 22 (1) | 0.415 | 3.274 |
| Sespe Creek | *M. brevipes* | SxB | 38 (2) | 0.409 | 3.384 |
| Sespe Creek | *M. fremontii* | SxF | 15 (1)<sup>†</sup> | 0.224<sup>†</sup> | 2.965<sup>†</sup> |
| Sespe Creek | Sespe Creek | SxS | 61 (6) | 0.757 | 1.757 |

**Bolded** values are significant in Tukey tests compared to both parental cross types.
<sup>†</sup>Measured SxF 'seeds' were indistinguishable from unfertilized ovules.

**Table S8.** Pollen viability and counts with sample sizes.

| Maternal species | Paternal species | Cross type | Average pollen viability (number of individuals) <sup>†</sup> | Pollen per unit area (number of individuals) |
| --- | --- | --- | --- | --- |
| *M. brevipes* | *M. brevipes* | BxB | 0.839 (13) | 55.6 (13) |
| *M. brevipes* | *M. fremontii* | BxF | **0.235 (12)** | 16.6 (13) |
| *M. fremontii* | *M. fremontii* | FxF | 0.922 (8) | 5.6 (10) |

<sup>†</sup>excluding flowers with <10 counted pollen grains

**Table S9.**
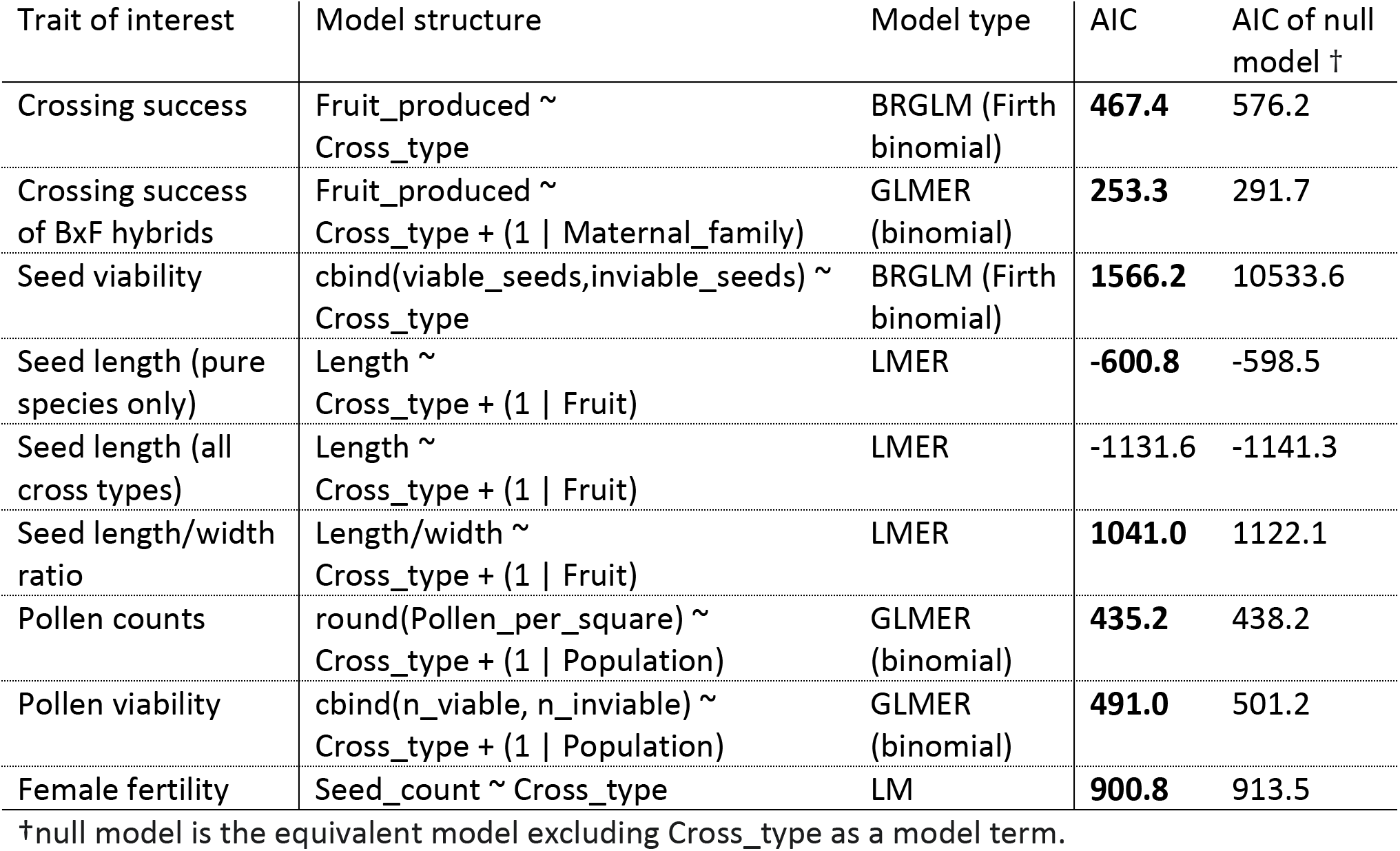
Statistical model structures and AIC values compared to null model equivalents.

## REFERENCES

Aagaard, J. E., George, R. D., Fishman, L., MacCoss, M. J., & Swanson, W. J. (2013). Selection on Plant Male Function Genes Identifies Candidates for Reproductive Isolation of Yellow Monkeyflowers. PLoS Genetics, 9(12). https://doi.org/10.1371/journal.pgen.1003965

Ascher, P. D., & Drewlow, L. W. (1975). The effect of prepollination injection with stigmatic exudate on interspecific pollen tube growth in Lilium longiflorum thunb. Styles. Plant Science Letters, 4(6), 401–405. https://doi.org/10.1016/0304-4211(75)90269-2

Baack, E., Melo, M. C., Rieseberg, L. H., & Ortiz-Barrientos, D. (2015). The origins of reproductive isolation in plants. New Phytologist, 207(4), 968–984. https://doi.org/10.1111/nph.13424

Baldwin, B. G., Goldman, D., Keil, D. J., Patterson, R., Rosatti, T. J., & Wilken, D. (Eds.). (2012). The Jepson Manual: Vascular Plants of California, Thoroughly Revised and Expanded (2nd ed.).

Barker, W. R. (Bill), Nesom, G. L., Beardsley, P. M., & Fraga, N. S. (2012). A taxonomic conspectus of phrymaceae: A narrowed circumscription for. *Phytoneuron*, May, 1–60.

Beardsley, P. M., Schoenig, S. E., Whittall, J. B., & Olmstead, R. G. (2004). Patterns of evolution in western North American Mimulus (Phrymaceae). American Journal of Botany, 91(3), 474–489. https://doi.org/10.3732/ajb.91.3.474

Behm, J. E., Ives, A. R., & Boughman, J. W. (2010). Breakdown in postmating isolation and the collapse of a species pair through hybridization. The American Naturalist, 175(1), 11–26. https://doi.org/10.1086/648559

Bernacchi, D., & Tanksley, S. D. (1997). An Interspecific Backcross of Lycopersicon esculentum × L. hirsutum: Linkage Analysis and a QTL Study of Sexual Compatibility Factors and Floral Traits. Genetics, 147(2), 861–877. https://doi.org/10.1093/genetics/147.2.861

Bickford, D., Lohman, D. J., Sodhi, N. S., Ng, P. K. L., Meier, R., Winker, K., Ingram, K. K., & Das, I. (2007). Cryptic species as a window on diversity and conservation. Trends in Ecology & Evolution, 22(3), 148–155. https://doi.org/10.1016/j.tree.2006.11.004

Bolger, A. M., Lohse, M., & Usadel, B. (2014). Trimmomatic: A flexible trimmer for Illumina sequence data. Bioinformatics, 30(15), 2114–2120. https://doi.org/10.1093/bioinformatics/btu170

Bowers, J. E. (1982). The plant ecology of inland dunes in western North America. Journal of Arid Environments, 5(3), 199–220. https://doi.org/10.1016/S0140-1963(18)31444-7

Brandvain, Y., & Haig, D. (2005). Divergent Mating Systems and Parental Conflict as a Barrier to Hybridization in Flowering Plants. The American Naturalist, 166(3), 330–338. https://doi.org/10.1086/432036

Brandvain, Y., Kenney, A. M., Flagel, L., Coop, G., & Sweigart, A. L. (2014). Speciation and Introgression between Mimulus nasutus and Mimulus guttatus. PLoS Genetics, 10(6), e1004410. https://doi.org/10.1371/journal.pgen.1004410

Broad Institute. (2019). Picard Toolkit, GitHub repository. https://broadinstitute.github.io/picard/

Christie, K., Fraser, L. S., & Lowry, D. B. (2022). The strength of reproductive isolating barriers in seed plants: Insights from studies quantifying premating and postmating reproductive barriers over the past 15 years. Evolution, evo.14565. https://doi.org/10.1111/evo.14565

Christie, K., & Strauss, S. Y. (2018). Along the speciation continuum: Quantifying intrinsic and extrinsic isolating barriers across five million years of evolutionary divergence in California jewelflowers. Evolution, 72(5), 1063–1079. https://doi.org/10.1111/evo.13477

Coughlan, J. M., & Matute, D. R. (2020). The importance of intrinsic postzygotic barriers throughout the speciation process: Intrinsic barriers throughout speciation. Philosophical Transactions of the Royal Society B: Biological Sciences, 375(1806). https://doi.org/10.1098/rstb.2019.0533

Coughlan, J. M., Wilson Brown, M., & Willis, J. H. (2020). Patterns of Hybrid Seed Inviability in the Mimulus guttatus sp. Complex Reveal a Potential Role of Parental Conflict in Reproductive Isolation. Current Biology, 30(1), 83–93.e5. https://doi.org/10.1016/j.cub.2019.11.023

Coyne, J. A., & Orr, H. A. (1989). Patterns of Speciation in Drosophila. Evolution, 43(2), 362–381.

Coyne, J. A., & Orr, H. A. (1997). Patterns of Speciation in Drosophila Revisited. Evolution, 51(1), 295. https://doi.org/10.2307/2410984

Crespi, B., & Nosil, P. (2013). Conflictual speciation: Species formation via genomic conflict. Trends in Ecology & Evolution, 28(1), 48–57. https://doi.org/10.1016/j.tree.2012.08.015

Curtu, A. L., Gailing, O., & Finkeldey, R. (2009). Patterns of contemporary hybridization inferred from paternity analysis in a four-oak-species forest. BMC Evolutionary Biology, 9(1), 284. https://doi.org/10.1186/1471-2148-9-284

Danecek, P., Bonfield, J. K., Liddle, J., Marshall, J., Ohan, V., Pollard, M. O., Whitwham, A., Keane, T., McCarthy, S. A., Davies, R. M., & Li, H. (2021). Twelve years of SAMtools and BCFtools. GigaScience, 10(2), giab008. https://doi.org/10.1093/gigascience/giab008

D’Angiolo, M., De Chiara, M., Yue, J.-X., Irizar, A., Stenberg, S., Persson, K., Llored, A., Barré, B., Schacherer, J., Marangoni, R., Gilson, E., Warringer, J., & Liti, G. (2020). A yeast living ancestor reveals the origin of genomic introgressions. Nature, 587(7834), Article 7834. https://doi.org/10.1038/s41586-020-2889-1

Darwin, C. (1859). On the origin of species by means of natural selection. John Murray. http://darwin-online.org.uk/converted/pdf/1861_OriginNY_F382.pdf

Ferris, K. G., Sexton, J. P., & Willis, J. H. (2014). Speciation on a local geographic scale: The evolution of a rare rock outcrop specialist in *Mimulus*. Philosophical Transactions of the Royal Society B: Biological Sciences, 369(1648), 20140001. https://doi.org/10.1098/rstb.2014.0001

Fishman, L. (2020). 96-well CTAB-chloroform DNA extraction. protocols.io. https://dx.doi.org/10.17504/protocols.io.bgv6jw9e

Fishman, L., Aagaard, J., & Tuthill, J. C. (2008). Toward the Evolutionary Genomics of Gametophytic Divergence: Patterns of Transmission Ratio Distortion in Monkeyflower (mimulus) Hybrids Reveal a Complex Genetic Basis for Conspecific Pollen Precedence. Evolution, 62(12), 2958–2970. https://doi.org/10.1111/j.1558-5646.2008.00475.x

Fishman, L., Sweigart, A. L., Kenney, A. M., & Campbell, S. (2014). Major quantitative trait loci control divergence in critical photoperiod for flowering between selfing and outcrossing species of monkeyflower (Mimulus). New Phytologist, 201(4), 1498–1507. https://doi.org/10.1111/nph.12618

Fishman, L., & Willis, J. H. (2006). A cytonuclear incompatibility causes anther sterility in Mimulus hybrids. Evolution, 60(7), 1372–1381. https://doi.org/10.1111/j.0014-3820.2006.tb01216.x

Fitak, R. R. (2021). OptM: Estimating the optimal number of migration edges on population trees using Treemix. Biology Methods and Protocols, 6(1), bpab017. https://doi.org/10.1093/biomethods/bpab017

Fujii, S., Tsuchimatsu, T., Kimura, Y., Ishida, S., Tangpranomkorn, S., Shimosato-Asano, H., Iwano, M., Furukawa, S., Itoyama, W., Wada, Y., Shimizu, K. K., & Takayama, S. (2019). A stigmatic gene confers interspecies incompatibility in the Brassicaceae. Nature Plants, 5(7), Article 7. https://doi.org/10.1038/s41477-019-0444-6

GBIF.org. (2022). GBIF Occurrence Download (12 September 2022). https://doi.org/10.15468/dl.j8dxeq

Gore, P., Potts, B., Volker, P., & Megalos, J. (1990). Unilateral Cross-Incompatibility in Eucalyptus: The Case of Hybridisation Between E. globulus and E. nitens. Australian Journal of Botany, 38(4), 383. https://doi.org/10.1071/BT9900383

Goulet, B. E., Roda, F., & Hopkins, R. (2017). Hybridization in plants: Old ideas, new techniques. Plant Physiology, 173(1), 65–78. https://doi.org/10.1104/pp.16.01340

Grant, A. L. (1924). A Monograph of the Genus Mimulus. Annals of the Missouri Botanical Garden, 11(2/3), 99–388. https://doi.org/10.2307/2394024

Grant, V. (1981). Plant Speciation. Columbia University Press.

Green, R. E., Krause, J., Briggs, A. W., Maricic, T., Stenzel, U., Kircher, M., Patterson, N., Li, H., Zhai, W., Fritz, M. H.-Y., Hansen, N. F., Durand, E. Y., Malaspinas, A.-S., Jensen, J. D., Marques-Bonet, T., Alkan, C., Prüfer, K., Meyer, M., Burbano, H. A., … Pääbo, S. (2010). A draft sequence of the Neandertal genome. *Science (New York*, N.Y.), 328(5979), 710– 722. https://doi.org/10.1126/science.1188021

Guerrero, R. F., Muir, C. D., Josway, S., & Moyle, L. C. (2017). Pervasive antagonistic interactions among hybrid incompatibility loci. PLOS Genetics, 13(6), e1006817. https://doi.org/10.1371/journal.pgen.1006817

Haig, D., & Westoby, M. (1991). Genomic imprinting in endosperm: Its effect on seed development in crosses between species, and between different ploidies of the same species, and its implications for the evolution of apomixis. *Philosophical Transactions - Royal Society of London*, B, 333, 1–13.

Hamlin, J. A. P., Hibbins, M. S., & Moyle, L. C. (2020). Assessing biological factors affecting postspeciation introgression. Evolution Letters, 4(2), 137–154. https://doi.org/10.1002/evl3.159

Hogenboom, N. G., Mather, K., Heslop-Harrison, J., & Lewis, D. (1975). Incompatibility and incongruity: Two different mechanisms for the non-functioning of intimate partner relationships. Proceedings of the Royal Society of London. Series B. Biological Sciences, 188(1092), 361–375. https://doi.org/10.1098/rspb.1975.0025

Kay, K. M., Woolhouse, S., Smith, B. A., Pope, N. S., & Rajakaruna, N. (2018). Sympatric serpentine endemic Monardella (Lamiaceae) species maintain habitat differences despite hybridization. Molecular Ecology, 27(9), 2302–2316. https://doi.org/10.1111/mec.14582

Kenney, A. M., & Sweigart, A. L. (2016). Reproductive isolation and introgression between sympatric Mimulus species. Molecular Ecology, 25(11), 2499–2517. https://doi.org/10.1111/mec.13630

Kleindorfer, S., O’Connor, J. A., Dudaniec, R. Y., Myers, S. A., Robertson, J., & Sulloway, F. J. (2014). Species collapse via hybridization in Darwin’s tree finches. The American Naturalist, 183(3), 325–341. https://doi.org/10.1086/674899

Koch, M. A., Haubold, B., & Mitchell-Olds, T. (2000). Comparative Evolutionary Analysis of Chalcone Synthase and Alcohol Dehydrogenase Loci in Arabidopsis, Arabis, and Related Genera (Brassicaceae). Molecular Biology and Evolution, 17(10), 1483–1498. https://doi.org/10.1093/oxfordjournals.molbev.a026248

Korunes, K. L., & Samuk, K. (2021). pixy: Unbiased estimation of nucleotide diversity and divergence in the presence of missing data. Molecular Ecology Resources, 21(4), 1359– 1368. https://doi.org/10.1111/1755-0998.13326

Kryvokhyzha, D. (n.d.). GATK: The best practice for genotype calling in a non-model organism. Dmytro Kryvokhyzha - Bioinformatics & Genomics Scientist. Retrieved August 8, 2022, from https://evodify.com/gatk-in-non-model-organism/

Kuboyama, T., Chung, C. S., & Takeda, G. (1994). The diversity of interspecific pollen-pistil incongruity in Nicotiana. Sexual Plant Reproduction, 7(4), 250–258. https://doi.org/10.1007/BF00232744

Lafon-Placette, C., Johannessen, I. M., Hornslien, K. S., Ali, M. F., Bjerkan, K. N., Bramsiepe, J., Glöckle, B. M., Rebernig, C. A., Brysting, A. K., Grini, P. E., & Köhler, C. (2017). Endosperm-based hybridization barriers explain the pattern of gene flow between Arabidopsis lyrata and Arabidopsis arenosa in Central Europe. Proceedings of the National Academy of Sciences, 114(6), E1027–E1035. https://doi.org/10.1073/pnas.1615123114

Lafon-Placette, C., & Köhler, C. (2016). Endosperm-based postzygotic hybridization barriers: Developmental mechanisms and evolutionary drivers. Molecular Ecology, 25(11), 2620– 2629. https://doi.org/10.1111/mec.13552

Le Gac, M., Hood, M. E., & Giraud, T. (2007). Evolution of Reproductive Isolation Within a Parasitic Fungal Species Complex. Evolution, 61(7), 1781–1787. https://doi.org/10.1111/j.1558-5646.2007.00144.x

Leishman, M., Wright, I., Moles, A., & Westoby, M. (2000). The Evolutionary Ecology of Seed Size. Seeds: The Ecology of Regeneration in Plant Communities, 2. https://doi.org/10.1079/9780851994321.0031

Li, H. (2018). Seqtk (1.3). https://github.com/lh3/seqtk

Li, H., & Durbin, R. (2009). Fast and accurate short read alignment with Burrows-Wheeler transform. Bioinformatics, 25(14), 1754–1760. https://doi.org/10.1093/bioinformatics/btp324

Lowry, D. B., Sobel, J. M., Angert, A. L., Ashman, T.-L., Baker, R. L., Blackman, B. K., Brandvain, Y., Byers, K. J. R. P., Cooley, A. M., Coughlan, J. M., Dudash, M. R., Fenster, C. B., Ferris, K. G., Fishman, L., Friedman, J., Grossenbacher, D. L., Holeski, L. M., Ivey, C. T., Kay, K. M., … Yuan, Y.-W. (2019). The case for the continued use of the genus name Mimulus for all monkeyflowers. TAXON, 68(4), 617–623. https://doi.org/10.1002/tax.12122

Lowry, D. B., & Willis, J. H. (2010). A widespread chromosomal inversion polymorphism contributes to a major life-history transition, local adaptation, and reproductive isolation. PLoS Biology, 8(9). https://doi.org/10.1371/journal.pbio.1000500

Malinsky, M., Matschiner, M., & Svardal, H. (2021). Dsuite—Fast D-statistics and related admixture evidence from VCF files. Molecular Ecology Resources, 21(2), 584–595. https://doi.org/10.1111/1755-0998.13265

Mallet, J., Beltrán, M., Neukirchen, W., & Linares, M. (2007). Natural hybridization in heliconiine butterflies: The species boundary as a continuum. BMC Evolutionary Biology, 7(1), 28. https://doi.org/10.1186/1471-2148-7-28

Malone, J. H., & Fontenot, B. E. (2008). Patterns of Reproductive Isolation in Toads. PLoS ONE, 3(12), e3900. https://doi.org/10.1371/journal.pone.0003900

Martin, S. H., & Van Belleghem, S. M. (2017). Exploring Evolutionary Relationships Across the Genome Using Topology Weighting. Genetics, 206(1), 429–438. https://doi.org/10.1534/genetics.116.194720

Matute, D. R., & Cooper, B. S. (2021). Comparative studies on speciation: 30 years since Coyne and Orr. Evolution, 1989, 1–15. https://doi.org/10.1111/evo.14181

Moyle, L. C., Olson, M. S., & Tiffin, P. (2004). Patterns of reproductive isolation in three angiosperm genera. Evolution, 58(6), 1195–1208. https://doi.org/10.1111/j.0014-3820.2004.tb01700.x

Moyle, L. C., & Payseur, B. A. (2009). Reproductive isolation grows on trees. Trends in Ecology & Evolution, 24(11), 591–598. https://doi.org/10.1016/j.tree.2009.05.010

Nelson, T. C., Stathos, A. M., Vanderpool, D. D., Finseth, F. R., Yuan, Y., & Fishman, L. (2021). Ancient and recent introgression shape the evolutionary history of pollinator adaptation and speciation in a model monkeyflower radiation (Mimulus section Erythranthe). PLoS Genetics, 17(2), e1009095. https://doi.org/10.1371/journal.pgen.1009095

Nesom, G. L. (2013). Taxonomic notes on Diplacus (Phrymaceae). Phytoneuron, 8.

Oneal, E., Willis, J. H., & Franks, R. G. (2016). Disruption of endosperm development is a major cause of hybrid seed inviability between *Mimulus guttatus* and *Mimulus nudatus*. New Phytologist, 210(3), 1107–1120. https://doi.org/10.1111/nph.13842

Pickrell, J. K., & Pritchard, J. K. (2012). Inference of Population Splits and Mixtures from Genome-Wide Allele Frequency Data. PLOS Genetics, 8(11), e1002967. https://doi.org/10.1371/journal.pgen.1002967

Presgraves, D. C. (2002). Patterns of Postzygotic Isolation in Lepidoptera. Evolution, 56(6), 1168–1183. https://doi.org/10.1111/j.0014-3820.2002.tb01430.x

Rabiee, M., Sayyari, E., & Mirarab, S. (2019). Multi-allele species reconstruction using ASTRAL. Molecular Phylogenetics and Evolution, 130, 286–296. https://doi.org/10.1016/j.ympev.2018.10.033

Ramsey, J., Bradshaw, H. D., & Schemske, D. W. (2003). Components of reproductive isolation between the monkeyflowers Mimulus lewisii and M. cardinalis (Phrymaceae). Evolution, 57(7), 1520–1534. https://doi.org/10.1111/j.0014-3820.2003.tb00360.x

Raunsgard, A., Opedal, Ø. H., Ekrem, R. K., Wright, J., Bolstad, G. H., Armbruster, W. S., & Pélabon, C. (2018). Intersexual conflict over seed size is stronger in more outcrossed populations of a mixed-mating plant. Proceedings of the National Academy of Sciences, 115(45), 11561–11566. https://doi.org/10.1073/pnas.1810979115

Rebernig, C. A., Lafon-Placette, C., Hatorangan, M. R., Slotte, T., & Köhler, C. (2015). Non- reciprocal Interspecies Hybridization Barriers in the Capsella Genus Are Established in the Endosperm. PLOS Genetics, 11(6), e1005295. https://doi.org/10.1371/journal.pgen.1005295

Roda, F., & Hopkins, R. (2019). Correlated evolution of self and interspecific incompatibility across the range of a Texas wildflower. New Phytologist, 221(1), 553–564. https://doi.org/10.1111/nph.15340

Roth, M., Florez-Rueda, A. M., & Städler, T. (2019). Differences in Effective Ploidy Drive Genome-Wide Endosperm Expression Polarization and Seed Failure in Wild Tomato Hybrids. Genetics, 212(1), 141–152. https://doi.org/10.1534/genetics.119.302056

Rushworth, C. A., Wardlaw, A. M., Ross-Ibarra, J., & Brandvain, Y. (2022). Conflict over fertilization underlies the transient evolution of reinforcement. PLOS Biology, 20(10), e3001814. https://doi.org/10.1371/journal.pbio.3001814

Sackton, T. (2022). Natural selection constrains neutral diversity across a wide range of species [R]. https://github.com/tsackton/linked-selection/blob/da62043544dce9c9342f0a393f41c63bec292fcf/misc_scripts/Identify_4D_Sites.pl (Original work published 2014)

Sandstedt, G. D., & Sweigart, A. L. (2022). Developmental evidence for parental conflict in driving Mimulus species barriers. New Phytologist, 236(4), 1545–1557. https://doi.org/10.1111/nph.18438

Sandstedt, G. D., Wu, C. A., & Sweigart, A. L. (2020). Evolution of multiple postzygotic barriers between species of the Mimulus tilingii complex. Evolution, 75, 600–613. https://doi.org/10.1111/evo.14105

Scascitelli, M., Whitney, K. D., Randell, R. A., King, M., Buerkle, C. A., & Rieseberg, L. H. (2010). Genome scan of hybridizing sunflowers from Texas ( Helianthus annuus and H. debilis ) reveals asymmetric patterns of introgression and small islands of genomic differentiation. Molecular Ecology, 19(3), 521–541. https://doi.org/10.1111/j.1365-294X.2009.04504.x

Schliep, K. P. (2011). phangorn: Phylogenetic analysis in R. Bioinformatics, 27(4), 592–593. https://doi.org/10.1093/bioinformatics/btq706

Schumer, M., Xu, C., Powell, D. L., Durvasula, A., Skov, L., Holland, C., Blazier, J. C., Sankararaman, S., Andolfatto, P., Gil, †, Rosenthal, G., & Przeworski, M. (2018). Natural selection interacts with recombination to shape the evolution of hybrid genomes. Science, 360(6389), 656–660.

Schwarzbach, A. E., Donovan, L. A., & Rieseberg, L. H. (2001). Transgressive character expression in a hybrid sunflower species. American Journal of Botany, 88(2), 270–277. https://doi.org/10.2307/2657018

Scopece, G., Musacchio, A., Widmer, A., & Cozzolino, S. (2007). Patterns of reproductive isolation in Mediterranean deceptive orchids. Evolution, 61(11), 2623–2642. https://doi.org/10.1111/j.1558-5646.2007.00231.x

Servedio, M. R., & Hermisson, J. (2020). The evolution of partial reproductive isolation as an adaptive optimum. Evolution, 74(1), 4–14. https://doi.org/10.1111/evo.13880

Skogsmyr, I., & Lankinen, Å. (2002). Sexual selection: An evolutionary force in plants? Biological Reviews, 77(4), 537–562. https://doi.org/10.1017/S1464793102005973

Sobel, J. M. (2010). Speciation in the western North American wildflower genus Mimulus [Ph.D., Michigan State University]. https://www.proquest.com/docview/873378456/abstract/950327CDE46D4690PQ/1

Sobel, J. M. (2014). Ecogeographic Isolation and Speciation in the Genus Mimulus. The American Naturalist, 184(5), 565–579. https://doi.org/10.1086/678235

Sobel, J. M., & Chen, G. F. (2014). Unification of methods for estimating the strength of reproductive isolation. Evolution, 68(5), 1511–1522. https://doi.org/10.1111/evo.12362

Stamatakis, A. (2014). RAxML version 8: A tool for phylogenetic analysis and post-analysis of large phylogenies. Bioinformatics, 30(9), 1312–1313. https://doi.org/10.1093/bioinformatics/btu033

Stankowski, S., Chase, M. A., Fuiten, A. M., Rodrigues, M. F., Ralph, P. L., & Streisfeld, M. A. (2019). Widespread selection and gene flow shape the genomic landscape during a radiation of monkeyflowers. PLOS Biology, 17(7), e3000391. https://doi.org/10.1371/journal.pbio.3000391

Stathos, A., & Fishman, L. (2014). Chromosomal rearrangements directly cause underdominant F1 pollen sterility in Mimulus lewisii–Mimulus cardinalis hybrids. Evolution, 68(11), 3109–3119. https://doi.org/10.1111/evo.12503

Streisfeld, M. A., & Kohn, J. R. (2005). Contrasting patterns of floral and molecular variation across a cline in Mimulus aurantiacus. Evolution, 59(12), 2548–2559. https://doi.org/10.1111/j.0014-3820.2005.tb00968.x

Streisfeld, M. A., & Rausher, M. D. (2009). Altered trans-Regulatory Control of Gene Expression in Multiple Anthocyanin Genes Contributes to Adaptive Flower Color Evolution in Mimulus aurantiacus. Molecular Biology and Evolution, 26(2), 433–444. https://doi.org/10.1093/molbev/msn268

Struck, T. H., Feder, J. L., Bendiksby, M., Birkeland, S., Cerca, J., Gusarov, V. I., Kistenich, S., Larsson, K.-H., Liow, L. H., Nowak, M. D., Stedje, B., Bachmann, L., & Dimitrov, D. (2018). Finding Evolutionary Processes Hidden in Cryptic Species. Trends in Ecology & Evolution, 33(3), 153–163. https://doi.org/10.1016/j.tree.2017.11.007

Sweigart, A. L., Fishman, L., & Willis, J. H. (2006). A simple genetic incompatibility causes hybrid male sterility in mimulus. Genetics, 172(4), 2465–2479. https://doi.org/10.1534/genetics.105.053686

Takeuchi, H., & Higashiyama, T. (2012). A Species-Specific Cluster of Defensin-Like Genes Encodes Diffusible Pollen Tube Attractants in Arabidopsis. PLOS Biology, 10(12), e1001449. https://doi.org/10.1371/journal.pbio.1001449

Tovar-Méndez, A., Kumar, A., Kondo, K., Ashford, A., Baek, Y. S., Welch, L., Bedinger, P. A., & McClure, B. A. (2014). Restoring pistil-side self-incompatibility factors recapitulates an interspecific reproductive barrier between tomato species. Plant Journal, 77(5), 727– 736. https://doi.org/10.1111/tpj.12424

Van der Auwera, G., & O’Connor, B. (2020). Genomics in the Cloud: Using Docker, GATK, and WDL in Terra (1st Edition). O’Reilly Media.

Wheeler, M. J., Franklin-Tong, V. E., & Franklin, F. C. H. (2001). The molecular and genetic basis of pollen–pistil interactions. New Phytologist, 151(3), 565–584. https://doi.org/10.1046/j.0028-646x.2001.00229.x

Williams, E., & Rouse, J. (1988). Disparate Style Lengths Contribute to Isolation of Species in Rhododendron. Australian Journal of Botany, 36(2), 183. https://doi.org/10.1071/BT9880183

Xiong, T., & Mallet, J. (2022). On the impermanence of species: The collapse of genetic incompatibilities in hybridizing populations. Evolution, 76(11), 2498–2512. https://doi.org/10.1111/evo.14626

Yuan, Y.-W. (2019). Monkeyflowers (Mimulus): New model for plant developmental genetics and evo-devo. New Phytologist, 222(2), 694–700. https://doi.org/10.1111/nph.15560

Yuan, Y.-W., Sagawa, J. M., Young, R. C., Christensen, B. J., & Bradshaw, H. D., Jr. (2013). Genetic Dissection of a Major Anthocyanin QTL Contributing to Pollinator-Mediated Reproductive Isolation Between Sister Species of Mimulus. Genetics, 194(1), 255–263. https://doi.org/10.1534/genetics.112.146852

Zuellig, M. P., & Sweigart, A. L. (2018). A two-locus hybrid incompatibility is widespread, polymorphic, and active in natural populations of Mimulus. Evolution, 72(11), 2394– 2405. https://doi.org/10.1111/evo.13596

